# Deciphering tea tree chloroplast and mitochondrial genomes of *Camellia sinensis* var. *assamica*

**DOI:** 10.1101/532853

**Authors:** Fen Zhang, Wei Li, Cheng-wen Gao, Li-zhi Gao

## Abstract

Tea is the most popular non-alcoholic caffeine-containing and the oldest beverage in the world. Despite its enormous industrial, cultural and medicinal values, the chloroplast (cp) and mitochondrial (mt) genomes are not available for *Camellia sinensis* var. *assamica*. In this study, we *de novo* assembled the cp genome sequence of *C. sinensis* var. *assamica* into a circular contig of 157,100 bp in length with an overall GC content of 37.29%, comprising a large single-copy region (LSC, 86,649 bp) and a small single-copy region (SSC, 18,285 bp) separated by a pair of inverted repeats (IRs, 26,083 bp). We annotated a total of 141 cp genes, of which 87 are protein-coding genes, 46 are tRNA genes, and eight are rRNA genes. We also *de novo* assembled the mt genome of *C. sinensis* var. *assamica* into two complete circular scaffolds (702,253 bp and 178,082 bp) with overall GC contents of 45.63% and 45.81%, respectively. We annotated a total of 71 mt genes, including 44 protein-coding genes, 24 tRNAs, and 3 rRNAs. Comparative analysis suggests repeat-rich nature of the mt genome compared to the cp genome, for example, with the characterization of 37,878 bp and 149 bp of long repeat sequences and 665 and 214 SSRs, respectively. We also detected 478 RNA-editing sites in 42 protein-coding mt genes, which are ∼4.4-fold more than 54 RNA-editing sites detected in 21 protein-coding cp genes. The high-quality cp and mt genomes of *C. sinensis* var. *assamica* presented in this study will become an invaluable resource for a range of genetic, functional, evolutionary and comparative genomic studies in tea tree and other *Camellia* species of the Theaceae family.

## INTRODUCTION

Tea is the most popular non-alcoholic caffeine-containing and the oldest beverage in the world since 3000 B. C. (Banerjee 1992; Mondal *et al.* 2004). The production of tea made from the young leaves of *Camellia sinensis* var. *sinensis* and *C. sinensis* var. *assamica*, together with ornamentally well-known camellias (e.g., *C. japonica, C. reticulata* and *C. sasanqua*) and worldwide renowned wooden crop *C. oleifera* (Ming and Bartholomew 2007) has made the genus *Camellia* possess huge economic values in Theaceae. Besides its industrial, cultural and medicinal values, botanists and evolutionary biologists have increasingly paid attention to this genus. As a result of frequent hybridization and polyploidization, *Camellia* is almost commonly regarded as one of the most taxonomically and phylogenetically difficult taxa in flowering plants (Huang *et al.* 2014). Thus, it has long been problematic for the taxonomic classification of the *Camellia* species based on the morphological characteristics (Lu *et al.* 2012). The chloroplast (cp) genomes are able to provide valuable information for taxonomic classification, tracing source populations (Mccauley *et al.* 1996; Small and Wendel 2004) and the reconstruction of phylogeny to resolve complex evolutionary relationships (Jansen *et al.* 2007; Moore *et al.* 2010; Parks *et al.* 2009) due to the conservation of genomic structure, maternal inheritance and a fairly low recombination rate. Genetically speaking, cp genomes are comparatively conserved than plant mitochondria (mt) genomes which are more heterogeneous in nature. It has long been acknowledged that mtDNA has the propensity to integrate DNA from various sources through intracellular and horizontal transfer (Schuster and Brennicke 1987; Stern and Lonsdale 1982; Vaughn *et al.* 1995). Partially due to these reasons, the mt genomes vary from ∼200 Kbp to ∼11.3 Mbp in some living organisms (Alverson *et al.* 2010; Sloan *et al.* 2012; Ward *et al.* 1981). The dynamic nature of mt genome structure has been recognized, and plant mt genomes can have a variety of different genomic configurations due to the recombination and differences in repeat content (Marechal and Brisson 2010; Palmer and Herbon 1988). These characteristics make the plant mt genome a fascinating genetic system to investigate questions related to evolutionary biology. Great efforts have been made to sequence the 13 representative *Camellia* chloroplast genomes using next-generation Illumina genome sequencing platform and obtain the first insight into global patterns of structural variation across the *Camellia* cp genomes (Huang *et al.* 2014). The reconstruction of phylogenetic relationships among these representative species of *Camellia* suggests that cp genomic resources are able to provide useful data to help to understand their evolutionary relationships and classify the ‘difficult taxa’. Recently, we decoded the first nuclear genome of *C. sinensis* var. *assamica* cv. *Yunkang10*, providing novel insights into genomic basis of tea flavors (Xia *et al.* 2017). Besides the absence of the *C. sinensis* var. *assamica* cp genome among 15 cp genomes that we have sequenced in this genus (Huang *et al.* 2014), none of mt genome is deciphered in the genus *Camellia*.

In this study, we filtered cpDNA and mtDNA reads from the WGS genome sequence project (Xia *et al.* 2017) and first *de novo* assembled the mt genome and cp genome of *C. sinensis* var. *assamica*. The information of both cp and mt genomes will help to obtain a comprehensive understanding of the taxonomy and evolution of the genus *Camellia*. These genome sequences will also facilitate the genetic modification of these economically important plants, for example, through chloroplast genetic engineering technologies.

## MATERIALS AND METHODS

### Plant materials, DNA extraction and genome sequencing

Young and healthy leaves of an individual plant of cultivar *Yunkang 10* of *C. sinensis* var. *assamica* were collected for genome sequencing in April, 2009, from Menghai County, Yunnan Province, China. Fresh leaves were harvested and immediately frozen in liquid nitrogen after collection, followed by the preservation at −80°C in the laboratory prior to DNA extraction. High-quality genomic DNA was extracted from leaves using a modified CTAB method (Porebski *et al.* 1997). RNase A and proteinase K were separately used to remove RNA and protein contamination. The quality and quantity of the isolated DNA were separately checked by electrophoresis on a 0.8% agarose gel and a NanoDrop D-1000 spectrophotometer (NanoDrop Technologies, Wilmington, DE). A total of eleven paired-end libraries, including four types of small-insert libraries (180 bp, 260 bp, 300 bp, 500 bp) and seven large-insert libraries (2 Kb, 3 Kb, 4 Kb, 5 Kb, 6 Kb, 8 Kb, 20 Kb), were prepared following the Illumina’s instructions, and sequenced using Illumina HiSeq2000 platform by following the standard Illumina protocols (Illumina, San Diego, CA). We totally generated ∼707.88 Gb (∼229.31×) of raw sequencing data (Xia *et al.* 2017). Further reads quality control filtering processes yielded a total of ∼492.15 Gb (∼159.43×) high-quality data retained and used for subsequent genome assembly.

### *De novo* chloroplast and mitochondria genome assemblies

The chloroplast reads were filtered from whole genome Illumina sequencing data of *C. sinensis* var. *assamica*, we mapped all the sequencing reads to the reference genomes (Huang *et al.* 2014) using bowtie2 (version 2.3.4.3) (Langmead *et al.* 2009). The mapped chloroplast reads were assembled using CLC Genomics Workbench v. 3.6.1 (CLC Inc., Rarhus, Denmark). For mitochondria genome assembly, the PE and MP sequencing reads were used separately. Briefly, we first performed *de novo* assembly with VELVET v1.2.08 (Zerbino and Birney 2008), which was previously described (Grewe *et al.* 2014; Zhu *et al.* 2014). Scaffolds were constructed using SSPACE v.3.0 (Boetzer *et al.* 2011). False connection was manually removed based on the coverage and distances of paired reads. Gaps between scaffolds were then filled with GapCloser (version 1.12) (Luo *et al.* 2012) using all pair-end reads. The completed chloroplast and mitochondria genomes are publicly available in GeneBank under accession numbers XXXXX and XXXXX.

### Genome annotation and visualization

The complete chloroplast genome of *C. sinensis* var. *assamica* was preliminarily annotated using the online program DOGMA (Dual Organellar Genome Annotator) (Wyman *et al.* 2004) followed by manual correction. MITOFY (Alverson *et al.* 2010) was used to characterize the complement of protein-coding and rRNA genes in the mitochondrial genome. A tRNA gene search was carried out using the tRNA scan-SE software(version 1.3.1) (Lowe and Eddy 1997). Circular genome maps were drawn with OrganellarGenomeDRAW (Lohse *et al.* 2007). Simple sequence repeats (SSRs) were identified and located using MISA (http://pgrc.ipk-gatersleben.de/misa/). All the annotated SSRs were classified by the size and copy number of their tandemly repeated: monomer (one nucleotide, n ⩾ 8), dimer (two nucleotides, n ⩾ 4), trimer (three nucleotides, n ⩾ 4), tetramer (four nucleotides, n ⩾ 3), pentamer (five nucleotides, n ⩾3), hexamer (six nucleotides, n ⩾ 3). Repeat sequences including forward and palindromic repeats, were also searched by REPuter (Kurtz *et al.* 2001) with the following parameters: minimal length 50 nt; mismatch 3 nt.

### Prediction of RNA-editing sites

Putative RNA editing sites in protein-coding genes were predicted using the PREP-cp and PREP-mt Web-based program (http://prep.unl.edu/) (Mower 2005; Mower 2009). To achieve a balanced trade-off between the number of false positive and false negative sites, the cutoff score (C-value) was set to 0.8 and 0.6, respectively (Chaw *et al.* 2008). All other parameters were set to default. Screening score value = 1.0 at editing sites are identified as credible RNA editing sites.

### Phylogenetic analyses

A total of thirteen conserved mt protein-coding genes among *C. sinensis* var. *assamica* and 14 other plant species (https://www.ncbi.nlm.nih.gov/genome/browse#!/organelles/) were individually aligned with ClustalW (Larkin *et al.* 2007), and then concatenated to construct a contiguous sequence in the order of *cob, cox1, cox2, cox3, nad1, nad2, nad3, nad4, nad4L, nad5, nad6, nad7* and *nad9*. The selected 14 species includes *Cycas taitungensis, Ginkgo biloba, Triticum aestivum, Oryza sativa, Sorghum bicolor, Zea mays, Gossypium arboretum, G. barbadense, Carica papaya, Vitis vinifera, Hevea brasiliensis, Bupleurum falcatum, Glycine max* and *Salvia miltiorrhiza*. The alignment file was used for the construction of Neighbor-Joining Tree at 1000 bootstrap replicates with MEGA 7.0.26 (Kumar *et al.* 2016). We employed the same method was used for phylogenetic analysis with cp genomes using the GTR+R+ I model under the maximum likelihood (ML) inference in MEGA v.7.0 (Kumar *et al.* 2016). Besides *C. sinensis* var. *assamica* cv. *Yunkang 10*, we selected cp genomes from the eighteen *Camelia* species (*C. oleifera, C. crapnelliana, C. szechuanensis, C. mairei, C. elongata, C. grandibracteata, C. leptophylla, C. petelotii, C. pubicosta, C. reticulata, C. azalea, C. japonica, C. cuspidata, C. danzaiensis, C. impressinervis, C. pitardii, C. yunnanensis* and *C. taliensis*) using *Apterosperm oblata* as outgroup.

### Data availability

Assembled mt and cp genome sequences and accompanying gene annotations of *C. sinensis* var. *assamica* have been deposited in the Genome Sequence Archive (Genomics, Proteomics & Bioinformatics 2017) in BIG Data Center (Nucleic Acids Res 2018), Beijing Institute of Genomics (BIG), Chinese Academy of Sciences, under accession numbers XXX and XXX that are publicly accessible at http://bigd.big.ac.cn/bioproject.

## RESULTS AND DISCUSSION

### Genome assembly and gene annotation of the *C. sinensis* var. *assamica* cp and mt genomes

The pair-end and mate-pair reads were used to assemble the *C. sinensis* var. *assamica* cp genome into a circular contig of 157,100 bp in length with an overall GC content of 37.29% (**Figure 1**; **Table 1**). It is a typical circular structure, including a large single-copy region (LSC, 86,649 bp) and a small single-copy region (SSC, 18,285 bp) separated by a pair of inverted repeats (IRs, 26,083 bp). A total of 141 genes were annotated, of which 87were protein-coding genes, 46 were tRNA genes and eight were rRNA genes. We obtained the two complete circular scaffolds (702,253 bp and 178,082 bp) of the *C. sinensis* var. *assamica* mt genome from the *de-novo* assembly of the filtered mitochondrial reads (**Figures 2-4**; **Table 2**). The two scaffolds of the mt genome had overall GC contents of 45.63% and 45.81%, respectively. We annotated a total of 71 genes, including 44 protein-coding genes, 24 tRNAs and 3 rRNAs (**Table 2**). Among 44 protein-coding genes, there were six genes with double copies, including *nad1, nad2, nad*9, *sdh3, atp9* and *rps19*. Of the tRNA genes, there were two copies each for *trnfM (Met), trnC (Cys)* and *trnP (Pro)* and three copies for *trnS (Ser)*.

**Table 1.**
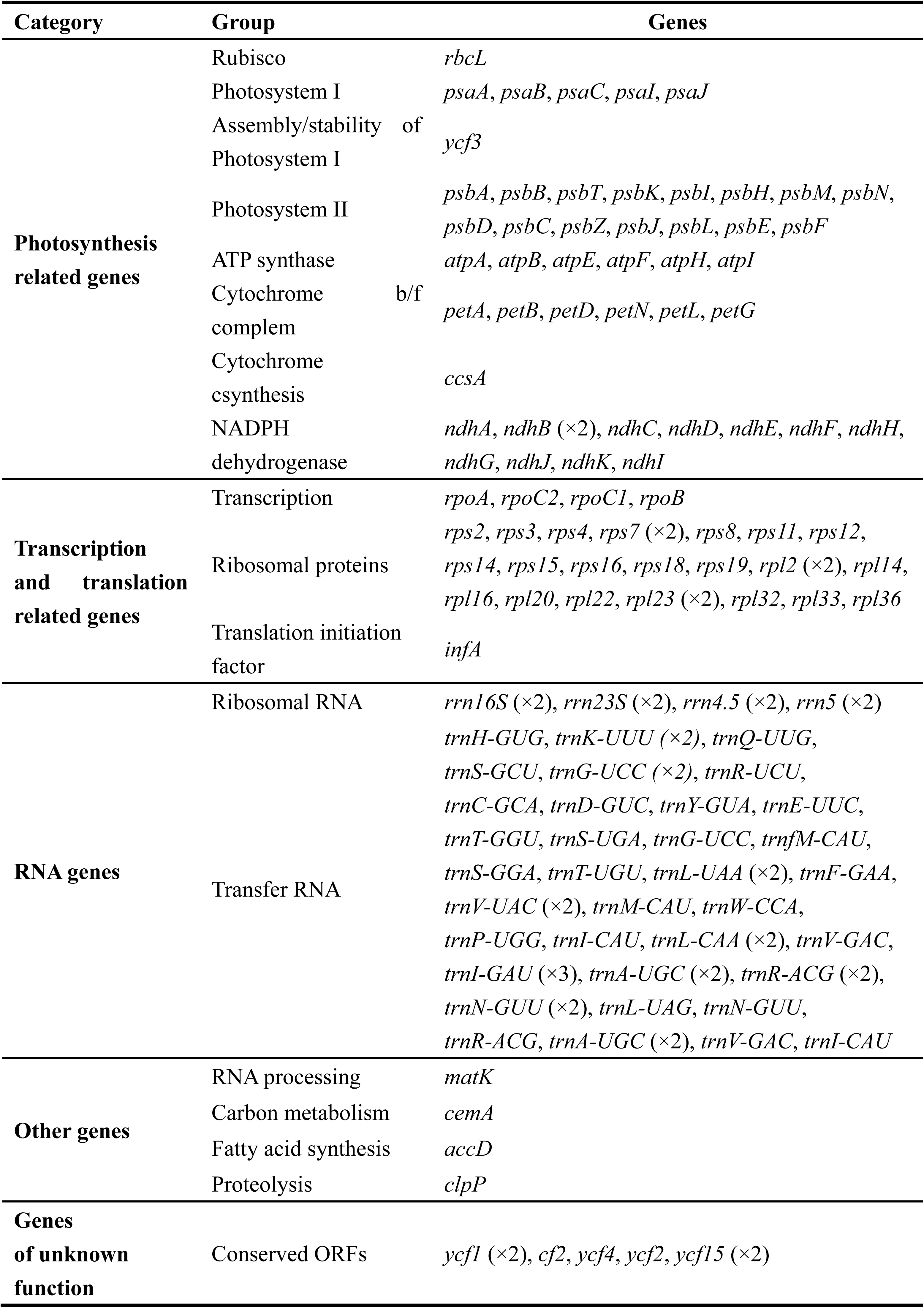
Gene annotation of the *C. sinensis* var. *assamica* cp genome.

**Table 2.**
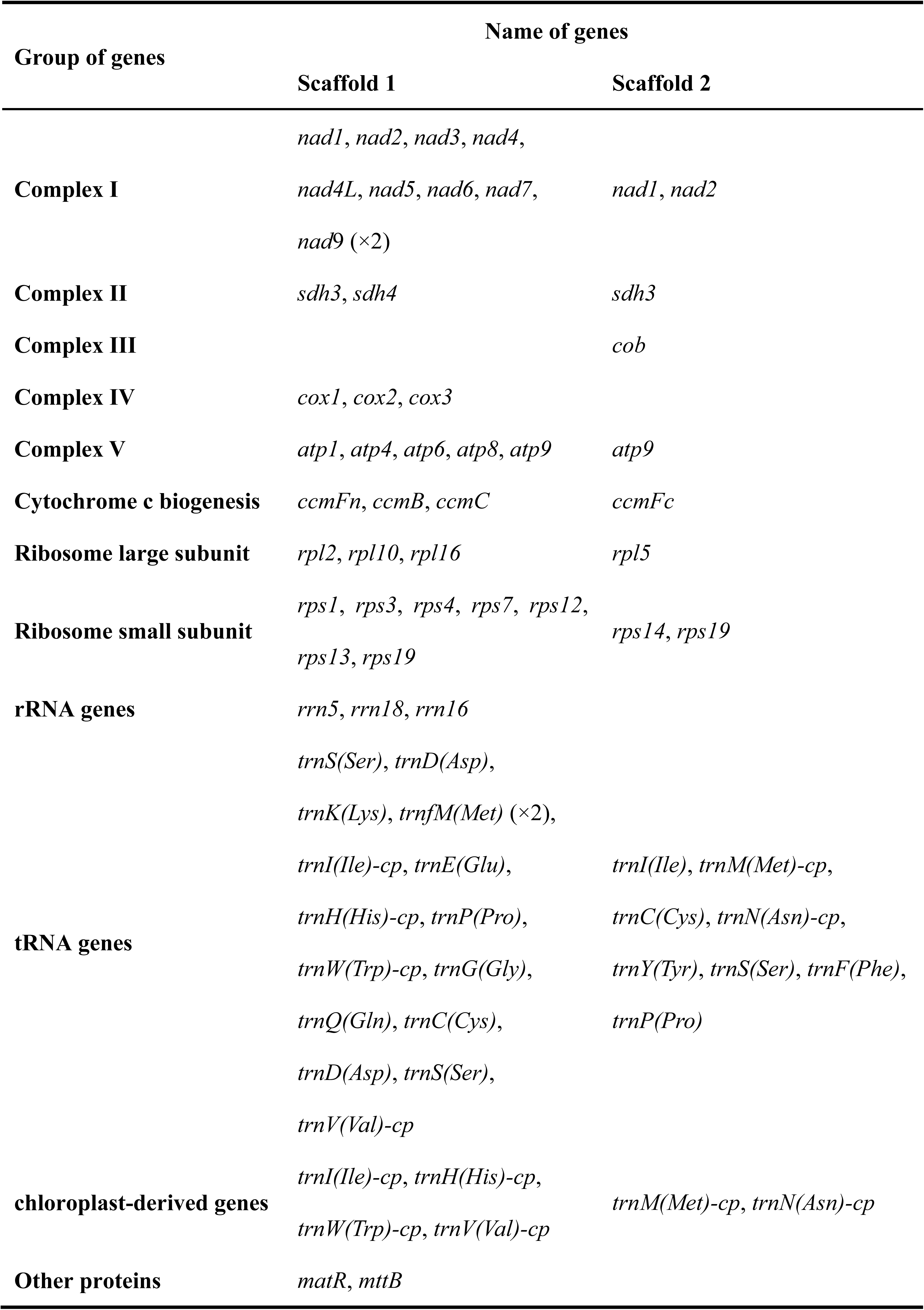
Gene content of of the *C. sinensis* var. *assamica* mt genome.

**Figure 1.**
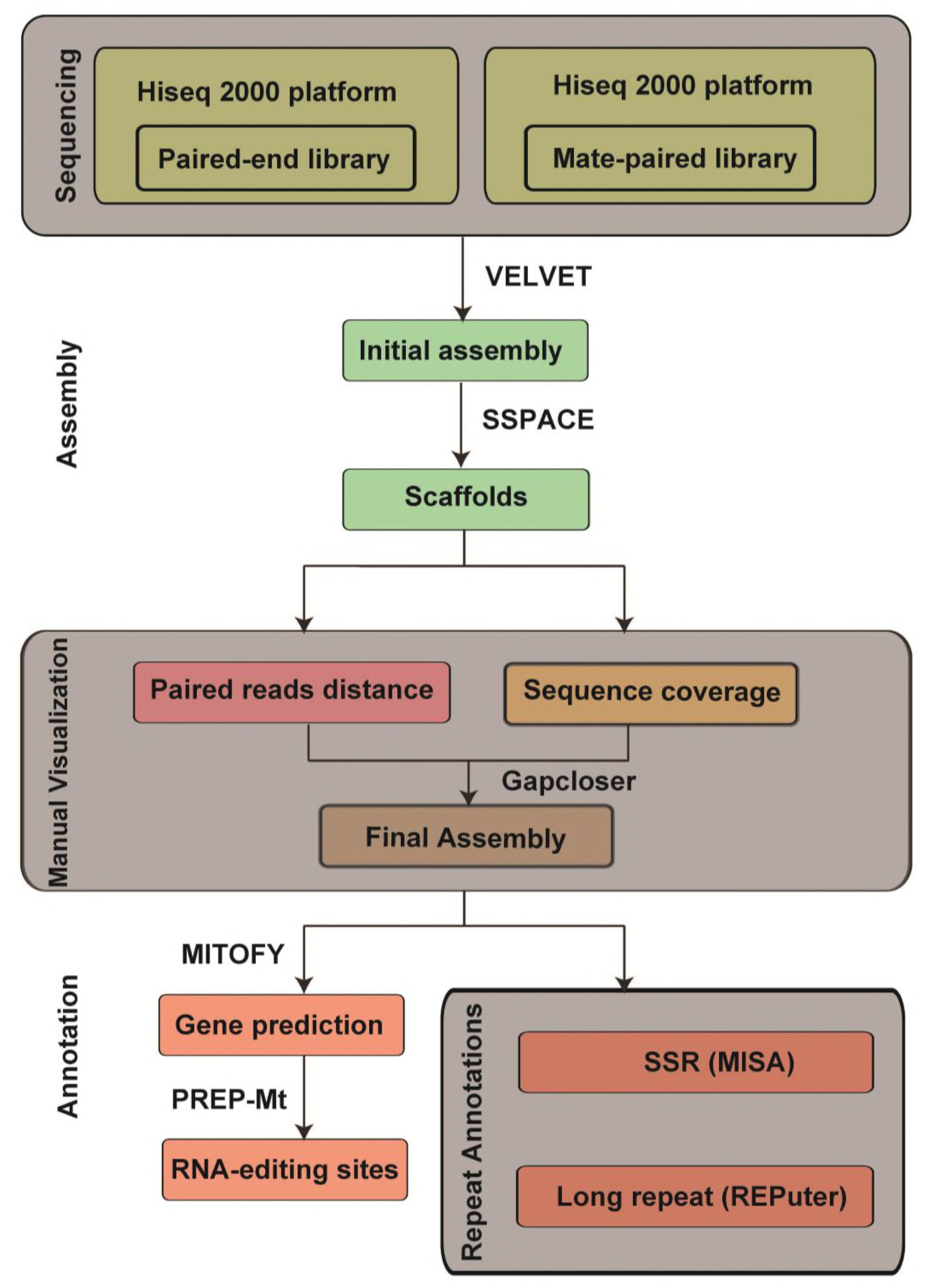
Workflow schema of the *Camellia sinensis* mitochondrial genome.

**Figure 2.**
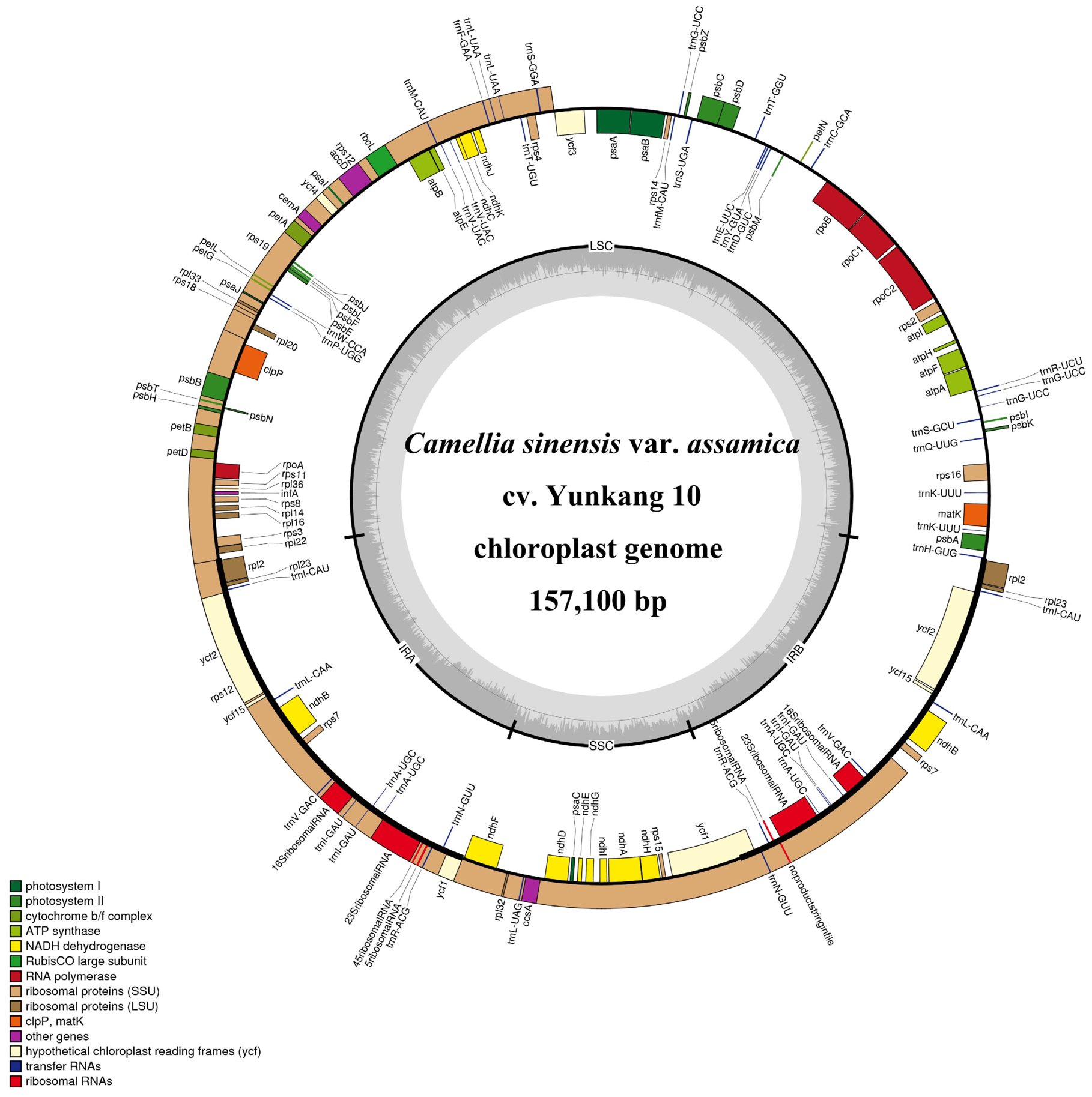
The assembly and annotation pipeline of the tea tree mitochondrial genome.

### Repeat contents of the *C. sinensis* var. *assamica* mt and cp genomes

A total of 214 SSRs were identified in cp genome with 74.42% of which were monomers, 19.07% of dimers, 0.47% of trimers, 4.65% of tetramers and 0.93% of hexamers (**Table 3**). There were no pentamers found in the cp genome. In mt genome, we discovered 665 SSRs distributed into monomers, dimers, trimers, pentamers, tetramers and hexamers with 31.53%, 45.35%, 4.95%, 15.17%, 2.70% and 0.15%, respectively (**Table 3**). Long repeat sequences (repeat unit > 50 bp) of forward and palindromic repeats were further annotated, resulting in 149 bp from 4 paired repeats in the cp genome (**Table 4**) and 37,878 bp from 58 paired repeats in the mt genome (**Table 5**). Our repeat content analyses indicate that the mt genome is more abundant in repeat sequences and more variable than the cp genome of *C. sinensis* var. *assamica* (**Table 5**).

**Table 3.** Statistics of SSR motifs in the *C. sinensis* var. *assamica* mt and cp genomes.

| SSR-Motif | mt Genome |  | cp Genome |  |
| --- | --- | --- | --- | --- |
|  | SSR Number | SSR % | SSR Number | SSR % |
| **Monomer** | 210 | 31.53 | 160 | 74.42 |
| **Dimer** | 302 | 45.35 | 41 | 19.07 |
| **Trimer** | 33 | 4.95 | 1 | 0.47 |
| **Tetramer** | 101 | 15.17 | 10 | 4.65 |
| **Pentamer** | 18 | 2.70 | 0 | 0.00 |
| **Hexamer** | 1 | 0.15 | 2 | 0.93 |

**Table 4.** Long repeats (repeat unit > 50 bp) in the *C. sinensis* var. *assamica* cp genome*.

| Repeat Length | Type | Start of Copy 1 | Start of Copy 2 |
| --- | --- | --- | --- |
| 56 | F | 93938 | 93956 |
| 56 | P | 93938 | 149737 |
| 56 | P | 93956 | 149755 |
| 56 | F | 149737 | 149755 |

\* P indicates palindromic repeats; F indicates forward repeats.
Overlapped repeats have been manually removed while calculating total length.

**Table 5.**
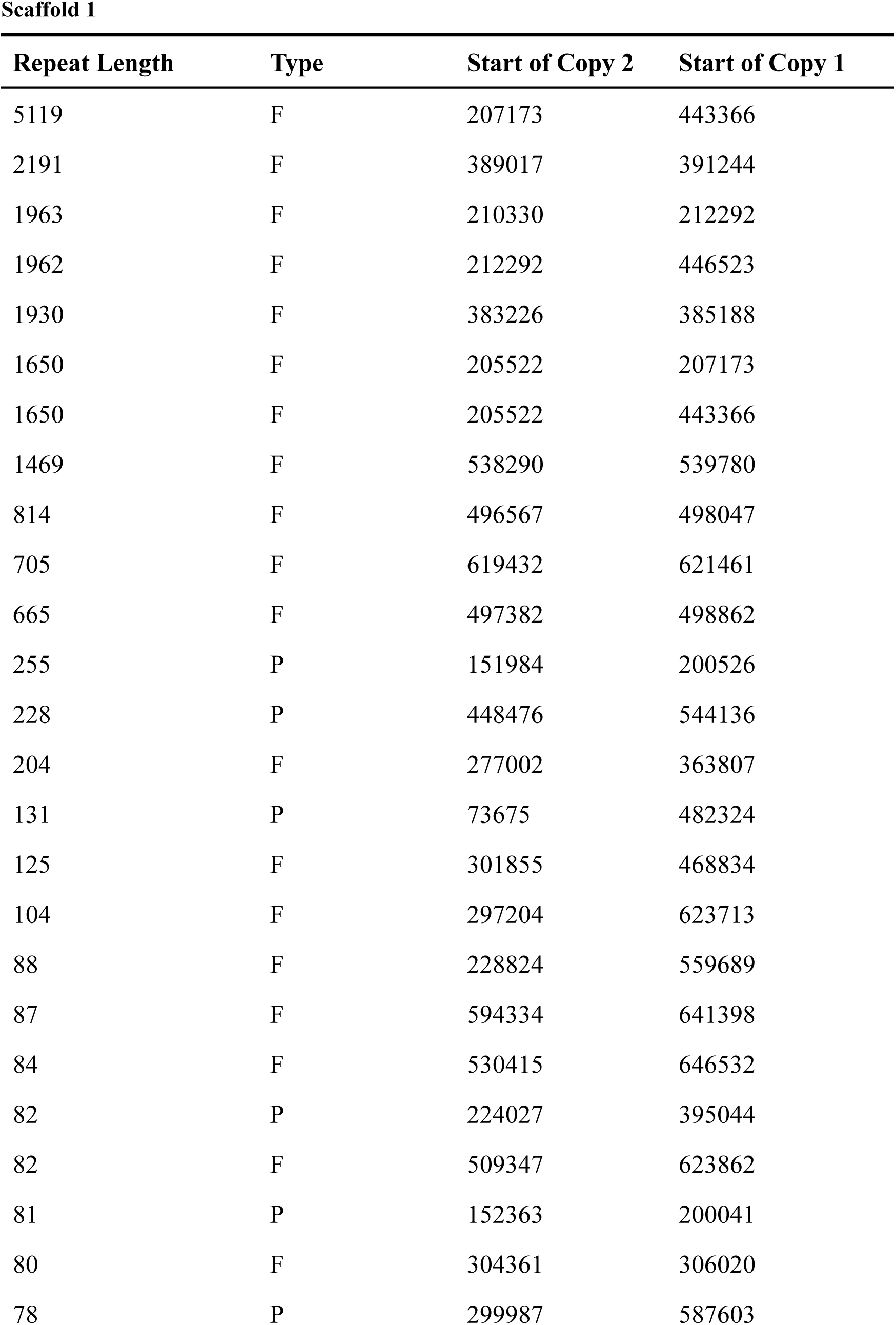
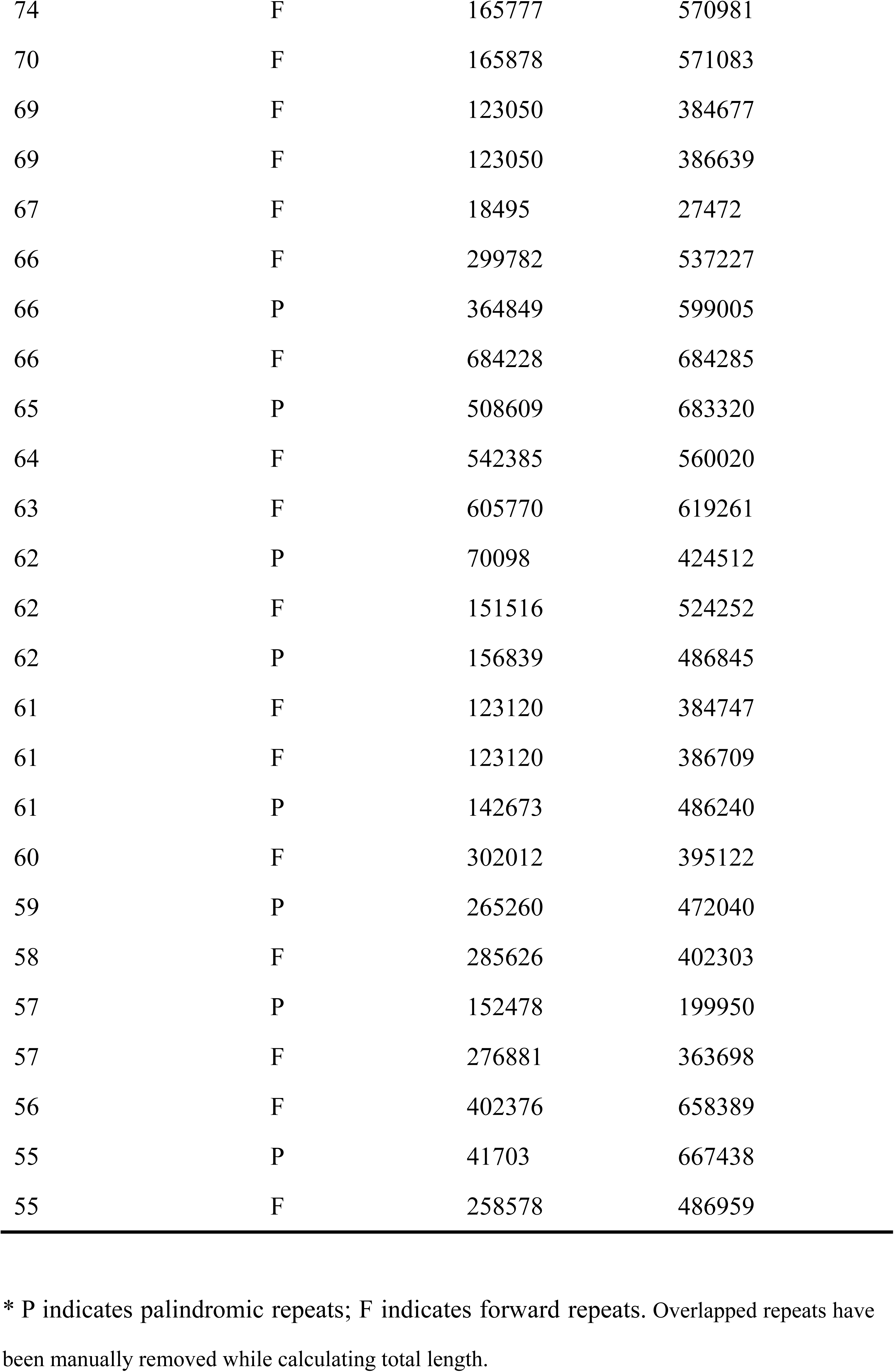
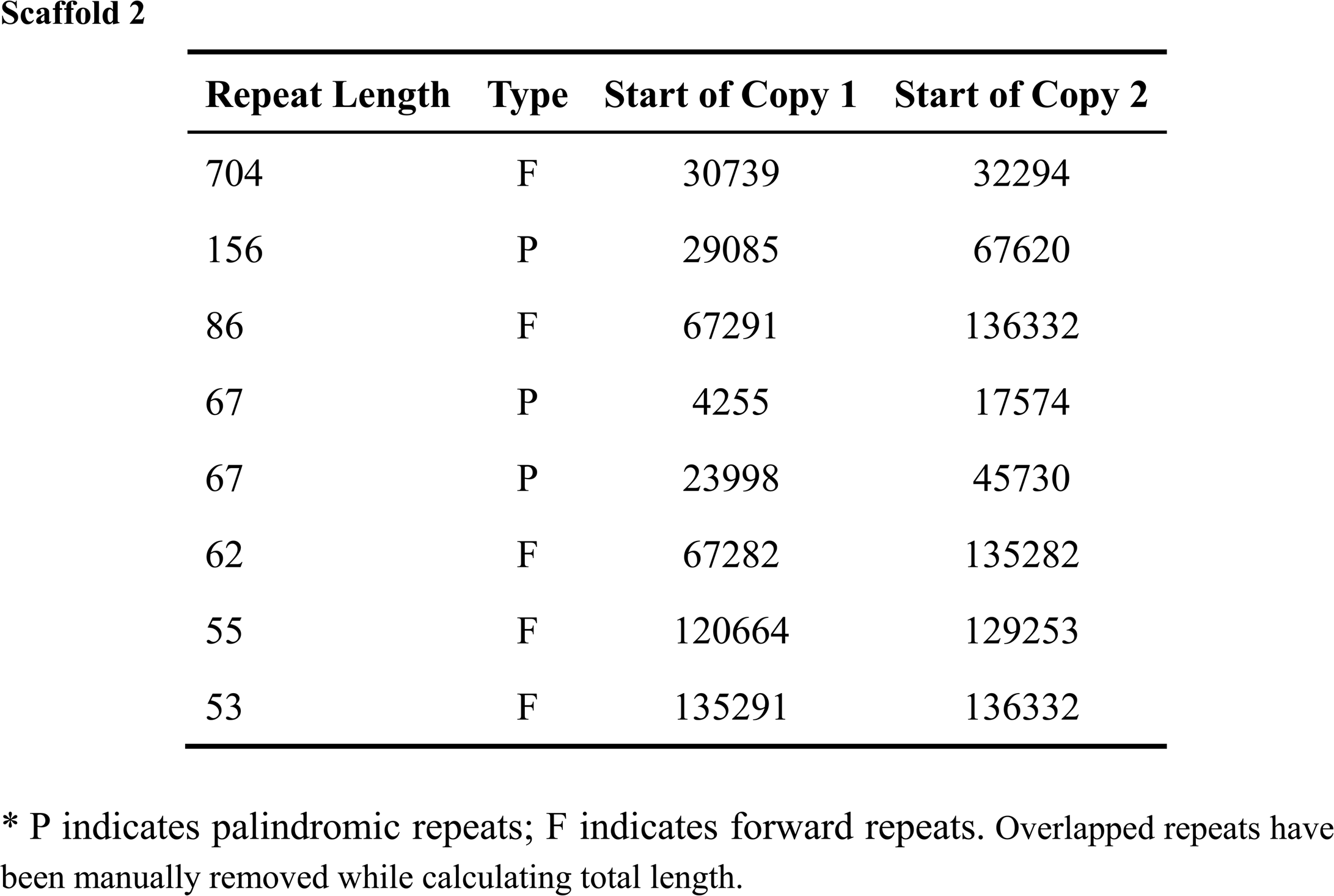
Long repeats (repeat unit > 50 bp) in the *C. sinensis* var. *assamica* mt genome*.

**Scaffold 1**
| Repeat Length | Type | Start of Copy 2 | Start of Copy 1 |
| --- | --- | --- | --- |
| 5119 | F | 207173 | 443366 |
| 2191 | F | 389017 | 391244 |
| 1963 | F | 210330 | 212292 |
| 1962 | F | 212292 | 446523 |
| 1930 | F | 383226 | 385188 |
| 1650 | F | 205522 | 207173 |
| 1650 | F | 205522 | 443366 |
| 1469 | F | 538290 | 539780 |
| 814 | F | 496567 | 498047 |
| 705 | F | 619432 | 621461 |
| 665 | F | 497382 | 498862 |
| 255 | P | 151984 | 200526 |
| 228 | P | 448476 | 544136 |
| 204 | F | 277002 | 363807 |
| 131 | P | 73675 | 482324 |
| 125 | F | 301855 | 468834 |
| 104 | F | 297204 | 623713 |
| 88 | F | 228824 | 559689 |
| 87 | F | 594334 | 641398 |
| 84 | F | 530415 | 646532 |
| 82 | P | 224027 | 395044 |
| 82 | F | 509347 | 623862 |
| 81 | P | 152363 | 200041 |
| 80 | F | 304361 | 306020 |
| 78 | P | 299987 | 587603 |
| 74 | F | 165777 | 570981 |
| 70 | F | 165878 | 571083 |
| 69 | F | 123050 | 384677 |
| 69 | F | 123050 | 386639 |
| 67 | F | 18495 | 27472 |
| 66 | F | 299782 | 537227 |
| 66 | P | 364849 | 599005 |
| 66 | F | 684228 | 684285 |
| 65 | P | 508609 | 683320 |
| 64 | F | 542385 | 560020 |
| 63 | F | 605770 | 619261 |
| 62 | P | 70098 | 424512 |
| 62 | F | 151516 | 524252 |
| 62 | P | 156839 | 486845 |
| 61 | F | 123120 | 384747 |
| 61 | F | 123120 | 386709 |
| 61 | P | 142673 | 486240 |
| 60 | F | 302012 | 395122 |
| 59 | P | 265260 | 472040 |
| 58 | F | 285626 | 402303 |
| 57 | P | 152478 | 199950 |
| 57 | F | 276881 | 363698 |
| 56 | F | 402376 | 658389 |
| 55 | P | 41703 | 667438 |
| 55 | F | 258578 | 486959 |

| **Repeat Length** | **Type** | **Start of Copy 1** | **Start of Copy 2** |
| --- | --- | --- | --- |
| 704 | F | 30739 | 32294 |
| 156 | P | 29085 | 67620 |
| 86 | F | 67291 | 136332 |
| 67 | P | 4255 | 17574 |
| 67 | P | 23998 | 45730 |
| 62 | F | 67282 | 135282 |
| 55 | F | 120664 | 129253 |
| 53 | F | 135291 | 136332 |

\* P indicates palindromic repeats; F indicates forward repeats. Overlapped repeats have been manually removed while calculating total length.

### Prediction of RNA-editing sites in the *C. sinensis* var. *assamica* cp and mt genomes

Almost all transcripts of protein encoding genes in the plant mitochondria are subject to RNA editing except the *T-urf13* gene (Ward and Levings 1991). Our results showed that the extent of RNA editing varied by gene for both cp and mt genomes of *C. sinensis* var. *assamica*. In the *C. sinensis* var. *assamica* cp genome, we detected 54 RNA-editing sites in 21 protein-coding genes, ranging from one editing site in *atpF, atpI, petB, psaI, psbE, psbF, rpoA, rps2* and *rps8* to 8 editing sites in *ndhB* (**Table 6**). In the *C. sinensis* var. *assamica* mt genome, we predicted 478 RNA-editing sites in 42 protein-coding genes; they varied from two editing site in *atp9* (of scaffold2), *sdh3* (of scaffold1 and scaffold2, respectively) and *rps14* (of scaffold2) to 35 editing sites in *ccmFn* (of scaffold1) (**Table 7-8**).

**Table 6.** Predicted RNA-editing sites in the *C. sinensis* var. *assamica* cp genome*. (i*The cutoff score (C-value) was set to 0.8.)

| No. | Gene | Nucleotide<br>Pos | AA<br>Pos | Effect | Score |
| --- | --- | --- | --- | --- | --- |
| 1 | *accD* | 64 | 22 | CGG (R) => TGG (W) | 1 |
| 2 | *accD* | 1469 | 490 | CCT (P) => CTT (L) | 1 |
| 3 | *atpA* | 791 | 264 | CCA (P) => CTA (L) | 1 |
| 4 | *atpA* | 914 | 305 | TCA (S) => TTA (L) | 1 |
| 5 | *atpF* | 92 | 31 | CCA (P) => CTA (L) | 0.86 |
| 6 | *atpI* | 134 | 45 | GCT (A) => GTT (V) | 1 |
| 7 | *matK* | 445 | 149 | CAC (H) => TAC (Y) | 1 |
| 8 | *matK* | 467 | 156 | TCG (S) => TTG (L) | 1 |
| 9 | *matK* | 631 | 211 | CAT (H) => TAT (Y) | 1 |
| 10 | *matK* | 1234 | 412 | CAT (H) => TAT (Y) | 1 |
| 11 | *ndhA* | 341 | 114 | TCA (S) => TTA (L) | 1 |
| 12 | *ndhA* | 566 | 189 | TCA (S) => TTA (L) | 1 |
| 13 | *ndhA* | 1028 | 343 | TCT (S) => TTT (F) | 1 |
| 14 | *ndhA* | 1073 | 358 | TCT (S) => TTT (F) | 1 |
| 15 | *ndhB* | 149 | 50 | TCA (S) => TTA (L) | 1 |
| 16 | *ndhB* | 467 | 156 | CCA (P) => CTA (L) | 1 |
| 17 | *ndhB* | 586 | 196 | CAT (H) => TAT (Y) | 1 |
| 18 | *ndhB* | 611 | 204 | TCA (S) => TTA (L) | 0.8 |
| 19 | *ndhB* | 737 | 246 | CCA (P) => CTA (L) | 1 |
| 20 | *ndhB* | 746 | 249 | TCT (S) => TTT (F) | 1 |
| 21 | *ndhB* | 830 | 277 | TCA (S) => TTA (L) | 1 |
| 22 | *ndhB* | 1481 | 494 | CCA (P) => CTA (L) | 1 |
| 23 | *ndhD* | 20 | 7 | ACG (T) => ATG (M) | 1 |
| 24 | *ndhD* | 401 | 134 | TCA (S) => TTA (L) | 1 |
| 25 | *ndhD* | 692 | 231 | TCA (S) => TTA (L) | 1 |
| 26 | *ndhD* | 896 | 299 | TCA (S) => TTA (L) | 1 |
| 27 | *ndhD* | 905 | 302 | CCT (P) => CTT (L) | 1 |
| 28 | *ndhD* | 1328 | 443 | TCA (S) => TTA (L) | 0.8 |
| 29 | *ndhF* | 205 | 69 | CAT (H) => TAT (Y) | 0.8 |
| 30 | *ndhF* | 290 | 97 | TCA (S) => TTA (L) | 1 |
| 31 | *ndhG* | 166 | 56 | CAT (H) => TAT (Y) | 0.8 |
| 32 | *ndhG* | 314 | 105 | ACA (T) => ATA (I) | 0.8 |
| 33 | *petB* | 641 | 214 | CCA (P) => CTA (L) | 1 |
| 34 | *psaI* | 80 | 27 | TCT (S) => TTT (F) | 0.86 |
| 35 | *psbE* | 214 | 72 | CCT (P) => TCT (S) | 1 |
| 36 | *psbF* | 77 | 26 | TCT (S) => TTT (F) | 1 |
| 37 | *rpoA* | 368 | 123 | TCG (S) => TTG (L) | 1 |
| 38 | *rpoB* | 338 | 113 | TCT (S) => TTT (F) | 1 |
| 39 | *rpoB* | 473 | 158 | TCA (S) => TTA (L) | 0.86 |
| 40 | *rpoB* | 551 | 184 | TCA (S) => TTA (L) | 1 |
| 41 | *rpoB* | 566 | 189 | TCG (S) => TTG (L) | 1 |
| 42 | *rpoB* | 973 | 325 | CTT (L) => TTT (F) | 0.86 |
| 43 | *rpoB* | 2000 | 667 | TCT (S) => TTT (F) | 1 |
| 44 | *rpoB* | 2336 | 779 | ACA (T) => ATA (I) | 1 |
| 45 | *rpoC1* | 41 | 14 | TCA (S) => TTA (L) | 1 |
| 46 | *rpoC1* | 1556 | 519 | TCG (S) => TTG (L) | 1 |
| 47 | *rpoC2* | 1505 | 502 | ACG (T) => ATG (M) | 0.86 |
| 48 | *rpoC2* | 2290 | 764 | CGG (R) => TGG (W) | 1 |
| 49 | *rpoC2* | 2726 | 909 | ACT (T) => ATT (I) | 1 |
| 50 | *rpoC2* | 3728 | 1243 | TCA (S) => TTA (L) | 0.86 |
| 51 | *rps2* | 248 | 83 | TCA (S) => TTA (L) | 1 |
| 52 | *rps8* | 182 | 61 | TCA (S) => TTA (L) | 0.86 |
| 53 | *rps14* | 80 | 27 | TCA (S) => TTA (L) | 1 |
| 54 | *rps14* | 149 | 50 | CCA (P) => CTA (L) | 1 |

**Table 7.** Predicted RNA-editing sites in Scaffold 1 of the *C. sinensis* var. *assamica* mt genome*.

| No. | Gene | Nucleotide Position | AA Pos | Effect | Score |
| --- | --- | --- | --- | --- | --- |
| 1 | *matR* | 32 | 11 | TCC (S) => TTC (F) | 0.62 |
| 2 | *matR* | 236 | 79 | TCC (S) => TTC (F) | 0.62 |
| 3 | *matR* | 326 | 109 | CCA (P) => CTA (L) | 1 |
| 4 | *matR* | 917 | 306 | TCA (S) => TTA (L) | 1 |
| 5 | *matR* | 1442 | 481 | GCC (A) => GTC (V) | 0.62 |
| 6 | *matR* | 1667 | 556 | TCC (S) => TTC (F) | 1 |
| 7 | *matR* | 1688 | 563 | CCT (P) => CTT (L) | 1 |
| 8 | *matR* | 1708 | 570 | CGC (R) => TGC (C) | 1 |
| 9 | *matR* | 1744 | 582 | CAC (H) => TAC (Y) | 1 |
| 10 | *matR* | 1775 | 592 | CCG (P) => CTG (L) | 1 |
| 11 | *matR* | 1814 | 605 | CCA (P) => CTA (L) | 0.88 |
| 12 | *matR* | 1832 | 611 | TCA (S) => TTA (L) | 0.88 |
| 13 | *ccmFn* | 38 | 13 | CCG (P) => CTG (L) | 1 |
| 14 | *ccmFn* | 98 | 33 | CCT (P) => CTT (L) | 1 |
| 15 | *ccmFn* | 137 | 46 | TCG (S) => TTG (L) | 1 |
| 16 | *ccmFn* | 142 | 48 | CGT (R) => TGT (C) | 1 |
| 17 | *ccmFn* | 151 | 51 | CCT (P) => TCT (S) | 0.83 |
| 18 | *ccmFn* | 248 | 83 | TCA (S) => TTA (L) | 1 |
| 19 | *ccmFn* | 256 | 86 | CGG (R) => TGG (W) | 1 |
| 20 | *ccmFn* | 283 | 95 | CTT (L) => TTT (F) | 0.83 |
| 21 | *ccmFn* | 334 | 112 | CAT (H) => TAT (Y) | 0.67 |
| 22 | *ccmFn* | 356 | 119 | TCC (S) => TTC (F) | 0.67 |
| 23 | *ccmFn* | 391 | 131 | CCT (P) => TCT (S) | 1 |
| 24 | *ccmFn* | 478 | 160 | CGT (R) => TGT (C) | 0.83 |
| 25 | *ccmFn* | 706 | 236 | CCT (P) => TTT (F) | 0.67 |
| 26 | *ccmFn* | 707 | 236 | CCT (P) => TTT (F) | 0.67 |
| 27 | *ccmFn* | 716 | 239 | TCA (S) => TTA (L) | 0.83 |
| 28 | *ccmFn* | 754 | 252 | CGT (R) => TGT (C) | 1 |
| 29 | *ccmFn* | 776 | 259 | TCA (S) => TTA (L) | 1 |
| 30 | *ccmFn* | 788 | 263 | CCA (P) => CTA (L) | 1 |
| 31 | *ccmFn* | 803 | 268 | TCA (S) => TTA (L) | 1 |
| 32 | *ccmFn* | 893 | 298 | GCG (A) => GTG (V) | 1 |
| 33 | *ccmFn* | 952 | 318 | CGC (R) => TGC (C) | 1 |
| 34 | *ccmFn* | 1270 | 424 | CGG (R) => TGG (W) | 1 |
| 35 | *ccmFn* | 1298 | 433 | CCA (P) => CTA (L) | 1 |
| 36 | *ccmFn* | 1315 | 439 | CAT (H) => TAT (Y) | 1 |
| 37 | *ccmFn* | 1330 | 444 | CGG (R) => TGG (W) | 1 |
| 38 | *ccmFn* | 1348 | 450 | CGG (R) => TGG (W) | 1 |
| 39 | *ccmFn* | 1381 | 461 | CGG (R) => TGG (W) | 1 |
| 40 | *ccmFn* | 1399 | 467 | CGT (R) => TGT (C) | 1 |
| 41 | *ccmFn* | 1442 | 481 | TCG (S) => TTG (L) | 1 |
| 42 | *ccmFn* | 1462 | 488 | CTT (L) => TTT (F) | 1 |
| 43 | *ccmFn* | 1466 | 489 | CCA (P) => CTA (L) | 1 |
| 44 | *ccmFn* | 1478 | 493 | TCA (S) => TTA (L) | 1 |
| 45 | *ccmFn* | 1487 | 496 | TCT (S) => TTT (F) | 1 |
| 46 | *ccmFn* | 1513 | 505 | CCC (P) => TCC (S) | 1 |
| 47 | *ccmFn* | 1561 | 521 | CGG (R) => TGG (W) | 0.67 |
| 48 | *nad5* | 155 | 52 | CCG (P) => CTG (L) | 1 |
| 49 | *nad5* | 238 | 80 | CCG (P) => TCG (S) | 0.8 |
| 50 | *nad5* | 269 | 90 | TCC (S) => TTC (F) | 0.7 |
| 51 | *nad5* | 355 | 119 | CCT (P) => TTT (F) | 1 |
| 52 | *nad5* | 356 | 119 | CCT (P) => TTT (F) | 1 |
| 53 | *nad5* | 371 | 124 | CCA (P) => CTA (L) | 0.9 |
| 54 | *nad5* | 395 | 132 | TCT (S) => TTT (F) | 0.9 |
| 55 | *nad5* | 503 | 168 | CCT (P) => CTT (L) | 1 |
| 56 | *nad5* | 536 | 179 | CCT (P) => CTT (L) | 1 |
| 57 | *nad5* | 626 | 209 | TCT (S) => TTT (F) | 0.9 |
| 58 | *nad5* | 628 | 210 | CGC (R) => TGC (C) | 0.9 |
| 59 | *nad5* | 673 | 225 | CTT (L) => TTT (F) | 0.9 |
| 60 | *nad5* | 710 | 237 | TCG (S) => TTG (L) | 1 |
| 61 | *nad5* | 722 | 241 | TCA (S) => TTA (L) | 1 |
| 62 | *nad5* | 832 | 278 | CCA (P) => TCA (S) | 0.9 |
| 63 | *nad5* | 872 | 291 | ACG (T) => ATG (M) | 1 |
| 64 | *nad5* | 1307 | 436 | TCA (S) => TTA (L) | 1 |
| 65 | *nad4* | 29 | 10 | TCC (S) => TTC (F) | 0.67 |
| 66 | *nad4* | 74 | 25 | ACT (T) => ATT (I) | 0.89 |
| 67 | *nad4* | 77 | 26 | CCT (P) => CTT (L) | 0.78 |
| 68 | *nad4* | 107 | 36 | CCG (P) => CTG (L) | 1 |
| 69 | *nad4* | 154 | 52 | CCC (P) => TCC (S) | 1 |
| 70 | *nad4* | 158 | 53 | CCT (P) => CTT (L) | 1 |
| 71 | *nad4* | 166 | 56 | CGG (R) => TGG (W) | 1 |
| 72 | *nad4* | 197 | 66 | TCT (S) => TTT (F) | 1 |
| 73 | *nad4* | 362 | 121 | ACA (T) => ATA (I) | 0.89 |
| 74 | *nad4* | 368 | 123 | TCT (S) => TTT (F) | 1 |
| 75 | *nad4* | 376 | 126 | CGT (R) => TGT (C) | 0.78 |
| 76 | *nad4* | 403 | 135 | CGC (R) => TGC (C) | 1 |
| 77 | *nad4* | 416 | 139 | CCT (P) => CTT (L) | 0.89 |
| 78 | *nad4* | 433 | 145 | CTT (L) => TTT (F) | 1 |
| 79 | *nad4* | 436 | 146 | CCC (P) => TTC (F) | 0.89 |
| 80 | *nad4* | 437 | 146 | CCC (P) => TTC (F) | 0.89 |
| 81 | *nad4* | 449 | 150 | CCA (P) => CTA (L) | 1 |
| 82 | *nad4* | 547 | 183 | CTC (L) => TTC (F) | 0.67 |
| 83 | *nad4* | 1336 | 446 | CAC (H) => TAC (Y) | 1 |
| 84 | *nad4* | 1352 | 451 | CCG (P) => CTG (L) | 1 |
| 85 | *nad4* | 1357 | 453 | CGC (R) => TGC (C) | 1 |
| 86 | *atp6* | 37 | 13 | CCA (P) => TCA (S) | 0.75 |
| 87 | *atp6* | 116 | 39 | TCA (S) => TTA (L) | 1 |
| 88 | *atp6* | 167 | 56 | CCG (P) => CTG (L) | 1 |
| 89 | *atp6* | 173 | 58 | CCG (P) => CTG (L) | 1 |
| 90 | *atp6* | 224 | 75 | TCC (S) => TTC (F) | 1 |
| 91 | *atp6* | 229 | 77 | CGC (R) => TGC (C) | 0.75 |
| 92 | *atp6* | 236 | 79 | TCG (S) => TTG (L) | 0.67 |
| 93 | *atp6* | 254 | 85 | TCG (S) => TTG (L) | 1 |
| 94 | *atp6* | 262 | 88 | CGT (R) => TGT (C) | 1 |
| 95 | *atp6* | 269 | 90 | CCC (P) => CTC (L) | 1 |
| 96 | *atp6* | 401 | 134 | TCA (S) => TTA (L) | 1 |
| 97 | *atp6* | 460 | 154 | CCT (P) => TCT (S) | 1 |
| 98 | *atp6* | 463 | 155 | CAT (H) => TAT (Y) | 1 |
| 99 | *atp6* | 485 | 162 | CCA (P) => CTA (L) | 1 |
| 100 | *atp6* | 527 | 176 | TCA (S) => TTA (L) | 1 |
| 101 | *atp6* | 548 | 183 | TCC (S) => TTC (F) | 1 |
| 102 | *atp6* | 635 | 212 | CCG (P) => CTG (L) | 1 |
| 103 | *atp6* | 656 | 219 | TCA (S) => TTA (L) | 1 |
| 104 | *atp6* | 664 | 222 | CAT (H) => TAT (Y) | 1 |
| 105 | *atp6* | 671 | 224 | TCT (S) => TTT (F) | 1 |
| 106 | *atp6* | 680 | 227 | TCA (S) => TTA (L) | 1 |
| 107 | *atp6* | 707 | 236 | ACA (T) => ATA (I) | 0.92 |
| 108 | *atp6* | 718 | 240 | CAA (Q) => TAA (X) | 1 |
| 109 | *mttB* | 58 | 20 | CAT (H) => TAT (Y) | 0.88 |
| 110 | *mttB* | 83 | 28 | TCG (S) => TTG (L) | 0.88 |
| 111 | *mttB* | 91 | 31 | CCA (P) => TCA (S) | 1 |
| 112 | *mttB* | 127 | 43 | CGT (R) => TGT (C) | 0.88 |
| 113 | *mttB* | 134 | 45 | CCA (P) => CTA (L) | 0.62 |
| 114 | *mttB* | 164 | 55 | TCC (S) => TTC (F) | 0.75 |
| 115 | *mttB* | 196 | 66 | CCG (P) => TCG (S) | 1 |
| 116 | *mttB* | 253 | 85 | CGT (R) => TGT (C) | 0.62 |
| 117 | *mttB* | 290 | 97 | TCT (S) => TTT (F) | 1 |
| 118 | *mttB* | 299 | 100 | TCG (S) => TTG (L) | 0.75 |
| 119 | *ccmB* | 28 | 10 | CAT (H) => TAT (Y) | 0.89 |
| 120 | *ccmB* | 43 | 15 | CCC (P) => TCC (S) | 0.67 |
| 121 | *ccmB* | 71 | 24 | CCA (P) => CTA (L) | 1 |
| 122 | *ccmB* | 80 | 27 | TCG (S) => TTG (L) | 1 |
| 123 | *ccmB* | 128 | 43 | TCA (S) => TTA (L) | 1 |
| 124 | *ccmB* | 137 | 46 | TCC (S) => TTC (F) | 1 |
| 125 | *ccmB* | 149 | 50 | CCG (P) => CTG (L) | 1 |
| 126 | *ccmB* | 154 | 52 | CGG (R) => TGG (W) | 1 |
| 127 | *ccmB* | 160 | 54 | CCT (P) => TCT (S) | 0.67 |
| 128 | *ccmB* | 164 | 55 | CCG (P) => CTG (L) | 0.89 |
| 129 | *ccmB* | 172 | 58 | CCT (P) => TCT (S) | 0.89 |
| 130 | *ccmB* | 179 | 60 | CCT (P) => CTT (L) | 1 |
| 131 | *ccmB* | 193 | 65 | CCT (P) => TTT (F) | 0.89 |
| 132 | *ccmB* | 194 | 65 | CCT (P) => TTT (F) | 0.89 |
| 133 | *ccmB* | 286 | 96 | CGG (R) => TGG (W) | 1 |
| 134 | *ccmB* | 304 | 102 | CGT (R) => TGT (C) | 0.78 |
| 135 | *ccmB* | 313 | 105 | CGT (R) => TGT (C) | 0.89 |
| 136 | *ccmB* | 338 | 113 | CCG (P) => CTG (L) | 1 |
| 137 | *ccmB* | 367 | 123 | CGG (R) => TGG (W) | 0.78 |
| 138 | *ccmB* | 424 | 142 | CGT (R) => TGT (C) | 0.89 |
| 139 | *ccmB* | 428 | 143 | TCG (S) => TTG (L) | 1 |
| 140 | *ccmB* | 467 | 156 | TCG (S) => TTG (L) | 0.89 |
| 141 | *ccmB* | 476 | 159 | CCA (P) => CTA (L) | 0.89 |
| 142 | *ccmB* | 485 | 162 | TCA (S) => TTA (L) | 1 |
| 143 | *ccmB* | 494 | 165 | TCA (S) => TTA (L) | 1 |
| 144 | *ccmB* | 503 | 168 | CCA (P) => CTA (L) | 1 |
| 145 | *ccmB* | 512 | 171 | TCT (S) => TTT (F) | 1 |
| 146 | *ccmB* | 514 | 172 | CGT (R) => TGT (C) | 1 |
| 147 | *ccmB* | 551 | 184 | TCA (S) => TTA (L) | 1 |
| 148 | *ccmB* | 554 | 185 | TCG (S) => TTG (L) | 0.89 |
| 149 | *ccmB* | 566 | 189 | TCC (S) => TTC (F) | 0.78 |
| 150 | *ccmB* | 569 | 190 | TCT (S) => TTT (F) | 0.78 |
| 151 | *ccmB* | 572 | 191 | CCG (P) => CTG (L) | 1 |
| 152 | *ccmB* | 596 | 199 | TCG (S) => TTG (L) | 0.89 |
| 153 | *rpl10* | 101 | 34 | TCG (S) => TTG (L) | 0.83 |
| 154 | *rpl10* | 239 | 80 | TCG (S) => TTG (L) | 0.83 |
| 155 | *rpl10* | 314 | 105 | TCA (S) => TTA (L) | 0.83 |
| 156 | *rps7* | 152 | 51 | CCA (P) => CTA (L) | 0.75 |
| 157 | *rps7* | 343 | 115 | CAC (H) => TAC (Y) | 0.62 |
| 158 | *rps7* | 368 | 123 | TCA (S) => TTA (L) | 0.88 |
| 159 | *atp1* | 1039 | 347 | CCC (P) => TCC (S) | 1 |
| 160 | *atp1* | 1064 | 355 | TCG (S) => TTG (L) | 1 |
| 161 | *atp1* | 1178 | 393 | TCA (S) => TTA (L) | 0.9 |
| 162 | *atp1* | 1216 | 406 | CTT (L) => TTT (F) | 1 |
| 163 | *atp1* | 1292 | 431 | CCG (P) => CTG (L) | 0.8 |
| 164 | *atp1* | 1415 | 472 | CCA (P) => CTA (L) | 1 |
| 165 | *atp1* | 1490 | 497 | CCA (P) => CTA (L) | 0.9 |
| 166 | *atp9* | 20 | 7 | TCA (S) => TTA (L) | 1 |
| 167 | *atp9* | 50 | 17 | TCA (S) => TTA (L) | 1 |
| 168 | *atp9* | 82 | 28 | CTT (L) => TTT (F) | 1 |
| 169 | *atp9* | 92 | 31 | TCG (S) => TTG (L) | 1 |
| 170 | *atp9* | 134 | 45 | TCA (S) => TTA (L) | 1 |
| 171 | *atp9* | 182 | 61 | TCG (S) => TTG (L) | 1 |
| 172 | *atp9* | 191 | 64 | CCA (P) => CTA (L) | 1 |
| 173 | *atp9* | 212 | 71 | TCA (S) => TTA (L) | 1 |
| 174 | *atp9* | 215 | 72 | TCC (S) => TTC (F) | 1 |
| 175 | *atp9* | 223 | 75 | CGA (R) => TGA (X) | 1 |
| 176 | *sdh3* | 67 | 23 | CCC (P) => TCC (S) | 1 |
| 177 | *sdh3* | 376 | 126 | CTC (L) => TTC (F) | 0.83 |
| 178 | *rpl16* | 79 | 27 | CAG (Q) => TAG (X) | 1 |
| 179 | *rpl16* | 227 | 76 | ACT (T) => ATT (I) | 1 |
| 180 | *rpl16* | 355 | 119 | CTC (L) => TTC (F) | 0.89 |
| 181 | *rpl16* | 524 | 175 | CCA (P) => CTA (L) | 1 |
| 182 | *rpl16* | 530 | 177 | TCG (S) => TTG (L) | 0.75 |
| 183 | *rps3* | 314 | 105 | CCA (P) => CTA (L) | 0.86 |
| 184 | *rps3* | 647 | 216 | CCG (P) => CTG (L) | 1 |
| 185 | *rps3* | 674 | 225 | CCG (P) => CTG (L) | 0.86 |
| 186 | *rps3* | 785 | 262 | TCA (S) => TTA (L) | 1 |
| 187 | *rps3* | 838 | 280 | CGT (R) => TGT (C) | 1 |
| 188 | *rps3* | 902 | 301 | TCA (S) => TTA (L) | 0.86 |
| 189 | *rps19* | 62 | 21 | TCG (S) => TTG (L) | 1 |
| 190 | *rps19* | 109 | 37 | CCT (P) => TTT (F) | 1 |
| 191 | *rps19* | 110 | 37 | CCT (P) => TTT (F) | 1 |
| 192 | *rpl2* | 215 | 72 | CCA (P) => CTA (L) | 0.75 |
| 193 | *rpl2* | 329 | 110 | CCA (P) => CTA (L) | 1 |
| 194 | *rpl2* | 494 | 165 | GCG (A) => GTG (V) | 0.67 |
| 195 | *rpl2* | 517 | 173 | CTC (L) => TTC (F) | 1 |
| 196 | *rpl2* | 550 | 184 | CCC (P) => TCC (S) | 1 |
| 197 | *atp8* | 47 | 16 | TCA (S) => TTA (L) | 1 |
| 198 | *atp8* | 58 | 20 | CTC (L) => TTC (F) | 1 |
| 199 | *atp8* | 452 | 151 | CCA (P) => CTA (L) | 0.75 |
| 200 | *cox3* | 289 | 97 | CTT (L) => TTT (F) | 0.92 |
| 201 | *cox3* | 304 | 102 | CGG (R) => TGG (W) | 1 |
| 202 | *cox3* | 311 | 104 | TCT (S) => TTT (F) | 0.92 |
| 203 | *cox3* | 314 | 105 | TCT (S) => TTT (F) | 0.92 |
| 204 | *cox3* | 419 | 140 | CCC (P) => CTC (L) | 1 |
| 205 | *cox3* | 422 | 141 | CCT (P) => CTT (L) | 0.92 |
| 206 | *cox3* | 512 | 171 | TCA (S) => TTA (L) | 0.75 |
| 207 | *cox3* | 653 | 218 | TCG (S) => TTG (L) | 1 |
| 208 | *cox3* | 754 | 252 | CGG (R) => TGG (W) | 0.92 |
| 209 | *cox3* | 764 | 255 | CCA (P) => CTA (L) | 0.92 |
| 210 | *sdh4* | 155 | 52 | CCA (P) => CTA (L) | 0.88 |
| 211 | *sdh4* | 203 | 68 | CCA (P) => CTA (L) | 0.75 |
| 212 | *sdh4* | 259 | 87 | CAT (H) => TAT (Y) | 0.88 |
| 213 | *cox1* | 155 | 52 | TCT (S) => TTT (F) | 1 |
| 214 | *cox1* | 167 | 56 | TCT (S) => TTT (F) | 1 |
| 215 | *cox1* | 265 | 89 | CCA (P) => TCA (S) | 1 |
| 216 | *cox1* | 356 | 119 | TCA (S) => TTA (L) | 1 |
| 217 | *cox1* | 365 | 122 | TCT (S) => TTT (F) | 1 |
| 218 | *cox1* | 428 | 143 | TCC (S) => TTC (F) | 1 |
| 219 | *cox1* | 464 | 155 | TCA (S) => TTA (L) | 1 |
| 220 | *cox1* | 503 | 168 | CCA (P) => CTA (L) | 1 |
| 221 | *cox1* | 581 | 194 | TCT (S) => TTT (F) | 1 |
| 222 | *cox1* | 628 | 210 | CGG (R) => TGG (W) | 1 |
| 223 | *cox1* | 659 | 220 | CCC (P) => CTC (L) | 1 |
| 224 | *cox1* | 674 | 225 | TCC (S) => TTC (F) | 1 |
| 225 | *cox1* | 758 | 253 | ACA (T) => ATA (I) | 1 |
| 226 | *cox1* | 773 | 258 | TCT (S) => TTT (F) | 1 |
| 227 | *cox1* | 950 | 317 | TCC (S) => TTC (F) | 1 |
| 228 | *cox1* | 1099 | 367 | CAC (H) => TAC (Y) | 1 |
| 229 | *cox1* | 1187 | 396 | CCG (P) => CTG (L) | 0.89 |
| 230 | *cox1* | 1318 | 440 | CGT (R) => TGT (C) | 0.78 |
| 231 | *cox1* | 1346 | 449 | TCA (S) => TTA (L) | 1 |
| 232 | *cox1* | 1402 | 468 | CCA (P) => TCA (S) | 1 |
| 233 | *cox1* | 1412 | 471 | TCG (S) => TTG (L) | 1 |
| 234 | *nad7* | 38 | 13 | TCG (S) => TTG (L) | 0.75 |
| 235 | *nad7* | 77 | 26 | TCA (S) => TTA (L) | 1 |
| 236 | *nad7* | 83 | 28 | TCA (S) => TTA (L) | 1 |
| 237 | *nad7* | 137 | 46 | TCA (S) => TTA (L) | 1 |
| 238 | *nad7* | 205 | 69 | CAT (H) => TAT (Y) | 1 |
| 239 | *nad7* | 212 | 71 | TCA (S) => TTA (L) | 1 |
| 240 | *nad7* | 277 | 93 | CGT (R) => TGT (C) | 1 |
| 241 | *nad7* | 296 | 99 | TCA (S) => TTA (L) | 0.88 |
| 242 | *nad7* | 305 | 102 | TCA (S) => TTA (L) | 1 |
| 243 | *nad7* | 344 | 115 | TCA (S) => TTA (L) | 1 |
| 244 | *nad7* | 494 | 165 | TCC (S) => TTC (F) | 1 |
| 245 | *nad7* | 539 | 180 | TCA (S) => TTA (L) | 0.88 |
| 246 | *nad7* | 812 | 271 | TCA (S) => TTA (L) | 0.88 |
| 247 | *nad7* | 859 | 287 | CCT (P) => TCT (S) | 0.88 |
| 248 | *nad7* | 943 | 315 | CGT (R) => TGT (C) | 1 |
| 249 | *nad7* | 965 | 322 | TCT (S) => TTT (F) | 1 |
| 250 | *nad7* | 989 | 330 | TCT (S) => TTT (F) | 1 |
| 251 | *nad7* | 1010 | 337 | CCA (P) => CTA (L) | 1 |
| 252 | *nad7* | 1052 | 351 | TCT (S) => TTT (F) | 1 |
| 253 | *nad9* | 428 | 143 | TCC (S) => TTC (F) | 0.73 |
| 254 | *nad9* | 506 | 169 | TCT (S) => TTT (F) | 0.75 |
| 255 | *nad9* | 527 | 176 | CCA (P) => CTA (L) | 0.92 |
| 256 | *nad9* | 581 | 194 | TCG (S) => TTG (L) | 0.92 |
| 257 | *nad9* | 604 | 202 | CAT (H) => TAT (Y) | 1 |
| 258 | *nad9* | 712 | 238 | CCG (P) => TCG (S) | 0.83 |
| 259 | *nad9* | 742 | 248 | CGG (R) => TGG (W) | 1 |
| 260 | *nad9* | 782 | 261 | TCC (S) => TTC (F) | 1 |
| 261 | *nad9* | 812 | 271 | TCA (S) => TTA (L) | 1 |
| 262 | *nad9* | 853 | 285 | CTT (L) => TTT (F) | 1 |
| 263 | *nad9* | 953 | 318 | TCT (S) => TTT (F) | 1 |
| 264 | *nad4L* | 11 | 4 | TCT (S) => TTT (F) | 1 |
| 265 | *nad4L* | 17 | 6 | TCA (S) => TTA (L) | 1 |
| 266 | *nad4L* | 25 | 9 | CGG (R) => TGG (W) | 1 |
| 267 | *nad4L* | 56 | 19 | CCT (P) => CTT (L) | 1 |
| 268 | *nad4L* | 65 | 22 | TCA (S) => TTA (L) | 1 |
| 269 | *nad4L* | 70 | 24 | CCA (P) => TCA (S) | 1 |
| 270 | *nad4L* | 80 | 27 | TCA (S) => TTA (L) | 1 |
| 271 | *nad4L* | 101 | 34 | TCG (S) => TTG (L) | 0.88 |
| 272 | *nad4L* | 128 | 43 | TCG (S) => TTG (L) | 1 |
| 273 | *nad4L* | 149 | 50 | TCA (S) => TTA (L) | 0.75 |
| 274 | *nad4L* | 158 | 53 | TCA (S) => TTA (L) | 0.88 |
| 275 | *nad4L* | 167 | 56 | CCA (P) => CTA (L) | 0.88 |
| 276 | *nad4L* | 200 | 67 | TCA (S) => TTA (L) | 1 |
| 277 | *nad4L* | 251 | 84 | TCT (S) => TTT (F) | 0.88 |
| 278 | *atp4* | 71 | 24 | TCA (S) => TTA (L) | 1 |
| 279 | *atp4* | 89 | 30 | TCA (S) => TTA (L) | 1 |
| 280 | *atp4* | 118 | 40 | CGT (R) => TGT (C) | 0.71 |
| 281 | *atp4* | 215 | 72 | TCG (S) => TTG (L) | 1 |
| 282 | *atp4* | 248 | 83 | CCT (P) => CTT (L) | 1 |
| 283 | *atp4* | 395 | 132 | TCA (S) => TTA (L) | 1 |
| 284 | *atp4* | 407 | 136 | CCA (P) => CTA (L) | 0.71 |
| 285 | *atp4* | 416 | 139 | ACT (T) => ATT (I) | 0.86 |
| 286 | *ccmC* | 76 | 26 | CGG (R) => TGG (W) | 0.78 |
| 287 | *ccmC* | 103 | 35 | CAT (H) => TAT (Y) | 1 |
| 288 | *ccmC* | 115 | 39 | CGG (R) => TGG (W) | 0.78 |
| 289 | *ccmC* | 133 | 45 | CTT (L) => TTT (F) | 0.67 |
| 290 | *ccmC* | 161 | 54 | CCG (P) => CTG (L) | 0.78 |
| 291 | *ccmC* | 179 | 60 | GCG (A) => GTG (V) | 0.78 |
| 292 | *ccmC* | 184 | 62 | CGG (R) => TGG (W) | 1 |
| 293 | *ccmC* | 299 | 100 | TCT (S) => TTT (F) | 1 |
| 294 | *ccmC* | 331 | 111 | CGG (R) => TGG (W) | 1 |
| 295 | *ccmC* | 395 | 132 | TCG (S) => TTG (L) | 1 |
| 296 | *ccmC* | 400 | 134 | CTT (L) => TTT (F) | 0.89 |
| 297 | *ccmC* | 421 | 141 | CGT (R) => TGT (C) | 0.78 |
| 298 | *ccmC* | 436 | 146 | CCT (P) => TCT (S) | 0.89 |
| 299 | *ccmC* | 446 | 149 | CCG (P) => CTG (L) | 0.78 |
| 300 | *ccmC* | 451 | 151 | CCT (P) => TCT (S) | 1 |
| 301 | *ccmC* | 458 | 153 | TCA (S) => TTA (L) | 0.78 |
| 302 | *ccmC* | 463 | 155 | CGT (R) => TGT (C) | 1 |
| 303 | *ccmC* | 467 | 156 | GCT (A) => GTT (V) | 0.78 |
| 304 | *ccmC* | 473 | 158 | CCG (P) => CTG (L) | 1 |
| 305 | *ccmC* | 497 | 166 | TCT (S) => TTT (F) | 1 |
| 306 | *ccmC* | 521 | 174 | TCG (S) => TTG (L) | 1 |
| 307 | *ccmC* | 548 | 183 | TCT (S) => TTT (F) | 1 |
| 308 | *ccmC* | 568 | 190 | CCT (P) => TCT (S) | 1 |
| 309 | *ccmC* | 575 | 192 | CCC (P) => CTC (L) | 1 |
| 310 | *ccmC* | 605 | 202 | TCC (S) => TTC (F) | 1 |
| 311 | *ccmC* | 608 | 203 | CCC (P) => CTC (L) | 0.89 |
| 312 | *ccmC* | 614 | 205 | TCA (S) => TTA (L) | 0.78 |
| 313 | *ccmC* | 619 | 207 | CGT (R) => TGT (C) | 0.78 |
| 314 | *ccmC* | 650 | 217 | CCT (P) => CTT (L) | 0.78 |
| 315 | *ccmC* | 656 | 219 | CCA (P) => CTA (L) | 0.89 |
| 316 | *ccmC* | 673 | 225 | CCT (P) => TCT (S) | 0.78 |
| 317 | *cox2* | 71 | 24 | TCT (S) => TTT (F) | 1 |
| 318 | *cox2* | 161 | 54 | TCA (S) => TTA (L) | 0.95 |
| 319 | *cox2* | 163 | 55 | CGG (R) => TGG (W) | 1 |
| 320 | *cox2* | 253 | 85 | CGG (R) => TGG (W) | 1 |
| 321 | *cox2* | 278 | 93 | CCG (P) => CTG (L) | 1 |
| 322 | *cox2* | 379 | 127 | CGG (R) => TGG (W) | 1 |
| 323 | *cox2* | 443 | 148 | ACG (T) => ATG (M) | 1 |
| 324 | *cox2* | 461 | 154 | CCA (P) => CTA (L) | 1 |
| 325 | *cox2* | 476 | 159 | TCA (S) => TTA (L) | 1 |
| 326 | *cox2* | 544 | 182 | CCT (P) => TCT (S) | 1 |
| 327 | *cox2* | 557 | 186 | CCT (P) => CTT (L) | 1 |
| 328 | *cox2* | 581 | 194 | TCA (S) => TTA (L) | 1 |
| 329 | *cox2* | 632 | 211 | TCG (S) => TTG (L) | 0.84 |
| 330 | *cox2* | 698 | 233 | ACG (T) => ATG (M) | 1 |
| 331 | *cox2* | 742 | 248 | CGG (R) => TGG (W) | 1 |
| 332 | *rps13* | 5 | 2 | TCA (S) => TTA (L) | 0.6 |
| 333 | *rps13* | 26 | 9 | TCA (S) => TTA (L) | 0.9 |
| 334 | *rps13* | 56 | 19 | TCA (S) => TTA (L) | 0.9 |
| 335 | *rps13* | 100 | 34 | CGT (R) => TGT (C) | 0.9 |
| 336 | *rps13* | 287 | 96 | TCG (S) => TTG (L) | 1 |
| 337 | *rps4* | 133 | 45 | CCG (P) => TCG (S) | 0.67 |
| 338 | *rps4* | 164 | 55 | TCA (S) => TTA (L) | 1 |
| 339 | *rps4* | 184 | 62 | CCC (P) => TCC (S) | 0.83 |
| 340 | *rps4* | 193 | 65 | CAT (H) => TAT (Y) | 1 |
| 341 | *rps4* | 257 | 86 | CCA (P) => CTA (L) | 1 |
| 342 | *rps4* | 266 | 89 | CCA (P) => CTA (L) | 0.83 |
| 343 | *rps4* | 278 | 93 | TCG (S) => TTG (L) | 0.67 |
| 344 | *rps4* | 290 | 97 | CCG (P) => CTG (L) | 0.83 |
| 345 | *rps4* | 335 | 112 | CCG (P) => CTG (L) | 1 |
| 346 | *rps4* | 482 | 161 | TCA (S) => TTA (L) | 1 |
| 347 | *rps4* | 914 | 305 | TCG (S) => TTG (L) | 0.83 |
| 348 | *rps4* | 925 | 309 | CAT (H) => TAT (Y) | 0.83 |
| 349 | *rps4* | 935 | 312 | CCA (P) => CTA (L) | 0.67 |
| 350 | *rps4* | 950 | 317 | TCT (S) => TTT (F) | 1 |
| 351 | *rps4* | 1001 | 334 | CCA (P) => CTA (L) | 0.83 |
| 352 | *rps4* | 1010 | 337 | CCT (P) => CTT (L) | 1 |
| 353 | *rps4* | 1015 | 339 | CGG (R) => TGG (W) | 1 |
| 354 | *nad1* | 8 | 3 | CCT (P) => CTT (L) | 0.9 |
| 355 | *nad1* | 65 | 22 | TCC (S) => TTC (F) | 1 |
| 356 | *nad1* | 100 | 34 | CCT (P) => TCT (S) | 0.9 |
| 357 | *nad1* | 149 | 50 | GCG (A) => GTG (V) | 0.9 |
| 358 | *nad1* | 209 | 70 | TCC (S) => TTC (F) | 1 |
| 359 | *nad1* | 308 | 103 | TCA (S) => TTA (L) | 1 |
| 360 | *nad1* | 434 | 145 | ACT (T) => ATT (I) | 1 |
| 361 | *nad6* | 7 | 3 | CTT (L) => TTT (F) | 1 |
| 362 | *nad6* | 83 | 28 | TCG (S) => TTG (L) | 1 |
| 363 | *nad6* | 88 | 30 | CCC (P) => TTC (F) | 0.7 |
| 364 | *nad6* | 89 | 30 | CCC (P) => TTC (F) | 0.7 |
| 365 | *nad6* | 95 | 32 | CCA (P) => CTA (L) | 1 |
| 366 | *nad6* | 103 | 35 | CGC (R) => TGC (C) | 1 |
| 367 | *nad6* | 161 | 54 | CCA (P) => CTA (L) | 1 |
| 368 | *nad6* | 169 | 57 | CAT (H) => TAT (Y) | 1 |
| 369 | *nad6* | 191 | 64 | TCA (S) => TTA (L) | 1 |
| 370 | *nad6* | 446 | 149 | TCC (S) => TTC (F) | 1 |
| 371 | *nad6* | 463 | 155 | CCT (P) => TCT (S) | 0.8 |
| 372 | *nad6* | 569 | 190 | TCT (S) => TTT (F) | 1 |
| 373 | *nad2* | 26 | 9 | TCC (S) => TTC (F) | 0.89 |
| 374 | *nad2* | 203 | 68 | TCT (S) => TTT (F) | 0.67 |
| 375 | *nad2* | 206 | 69 | TCC (S) => TTC (F) | 1 |
| 376 | *nad2* | 230 | 77 | TCT (S) => TTT (F) | 1 |
| 377 | *nad2* | 236 | 79 | TCC (S) => TTC (F) | 0.67 |
| 378 | *nad2* | 251 | 84 | CCA (P) => CTA (L) | 1 |
| 379 | *nad2* | 262 | 88 | CGC (R) => TGC (C) | 1 |
| 380 | *nad2* | 289 | 97 | CAT (H) => TAT (Y) | 1 |
| 381 | *nad2* | 296 | 99 | TCA (S) => TTA (L) | 1 |
| 382 | *nad2* | 323 | 108 | CCT (P) => CTT (L) | 1 |
| 383 | *nad2* | 392 | 131 | TCG (S) => TTG (L) | 1 |
| 384 | *rps12* | 71 | 24 | TCG (S) => TTG (L) | 0.94 |
| 385 | *rps12* | 100 | 34 | CGC (R) => TGC (C) | 1 |
| 386 | *rps12* | 104 | 35 | CCG (P) => CTG (L) | 1 |
| 387 | *rps12* | 196 | 66 | CAC (H) => TAC (Y) | 0.94 |
| 388 | *rps12* | 221 | 74 | TCG (S) => TTG (L) | 0.88 |
| 389 | *rps12* | 269 | 90 | TCG (S) => TTG (L) | 0.94 |
| 390 | *rps12* | 284 | 95 | TCC (S) => TTC (F) | 0.76 |
| 391 | *nad3* | 5 | 2 | TCA (S) => TTA (L) | 0.79 |
| 392 | *nad3* | 44 | 15 | CCG (P) => CTG (L) | 1 |
| 393 | *nad3* | 62 | 21 | CCA (P) => CTA (L) | 0.95 |
| 394 | *nad3* | 80 | 27 | CCA (P) => CTA (L) | 1 |
| 395 | *nad3* | 146 | 49 | TCC (S) => TTC (F) | 1 |
| 396 | *nad3* | 208 | 70 | CCT (P) => TTT (F) | 0.95 |
| 397 | *nad3* | 209 | 70 | CCT (P) => TTT (F) | 0.95 |
| 398 | *nad3* | 215 | 72 | CCG (P) => CTG (L) | 1 |
| 399 | *nad3* | 230 | 77 | TCC (S) => TTC (F) | 0.86 |
| 400 | *nad3* | 247 | 83 | CCT (P) => TCT (S) | 1 |
| 401 | *nad3* | 251 | 84 | CCC (P) => CTC (L) | 0.91 |
| 402 | *nad3* | 266 | 89 | CCG (P) => CTG (L) | 1 |
| 403 | *nad3* | 275 | 92 | TCT (S) => TTT (F) | 1 |
| 404 | *nad3* | 317 | 106 | TCT (S) => TTT (F) | 0.95 |
| 405 | *nad3* | 344 | 115 | TCG (S) => TTG (L) | 1 |
| 406 | *nad3* | 349 | 117 | CGG (R) => TGG (W) | 1 |
| 407 | *rps1* | 23 | 8 | CCT (P) => CTT (L) | 0.67 |
| 408 | *rps1* | 56 | 19 | CCT (P) => CTT (L) | 0.67 |
| 409 | *rps1* | 380 | 127 | TCA (S) => TTA (L) | 0.67 |

\*The cutoff score (C-value) was set to 0.6.

### Phylogenetic relationships based on cp and mt genomes

To further determine the phylogenetic position of *C. sinensis* var. *assamica* we performed phylogenomic analysis of 20 complete cp genomes using *Apterosperm oblata* as outgroup. Our results showed that *C. sinensis* var. *assamica* was grouped with *C. grandibracteata* with 100% bootstrap support (**Figure 5**). We further examined phylogenetic relationships based on the thirteen conserved mt protein-coding genes between *C. sinensis* var. *assamica* and 14 other plant species. Our results showed that *C. sinensis* var. *assamica* is clearly grouped with other dicots that were separated from monocots of the angiosperms while the two gymnosperms (*Cycas taitungensis* and *Ginkgo biloba*) were formed the basal clade (**Figure 6**).

### Conclusions

We *de novo* assembled both cp and mt *C. sinensis* var. *assamica* genomes, of which the mt genome is the first reported in the Theaceae family. A large set of annotated cp and mt genes with known homology will aid further gene ontology and functional genomic analyses. There is no doubt that cp genome structure and organization is much more conserved than mt and nuclear genomes of *C. sinensis* var. *assamica*. The genome information serves as a powerful tool to better understand the taxonomy, phylogeny and evolution in the *Camellia* genus. Taken together, the high-quality cp and mt genomes of *C. sinensis* var. *assamica* presented in this study will become an invaluable genomic resource for a range of functional, evolutionary and comparative genomic studies in tea tree and other plant species of the Theaceae family.

## ACKNOWLEDGEMENTS

We would thank Yunnan Tea Research Institute for providing tea plant materials in this study. We are grateful An-dan Zhu for technical support and anonymous reviewers for valuable comments on the manuscript. This work was supported by the Project of Innovation Team of Yunnan Province and Ten Thousands Talents Program of China (to L. Z. Gao).

## ADDITIONAL INFORMATION

Competing financial interests: The authors declare no competing financial interests.

---

**Figure 1. Genome map of *C. sinensis* var. *assamica* cv. *Yunkang 10***. Genes lying outside of the outer circle are transcribed in the clockwise direction whereas genes inside are transcribed in the counterclockwise direction. Genes belonging to different functional groups are color-coded. Area dashed darker gray in the inner circle indicates GC content while the lighter gray corresponds to AT content of the genome.

**Figure 3.**
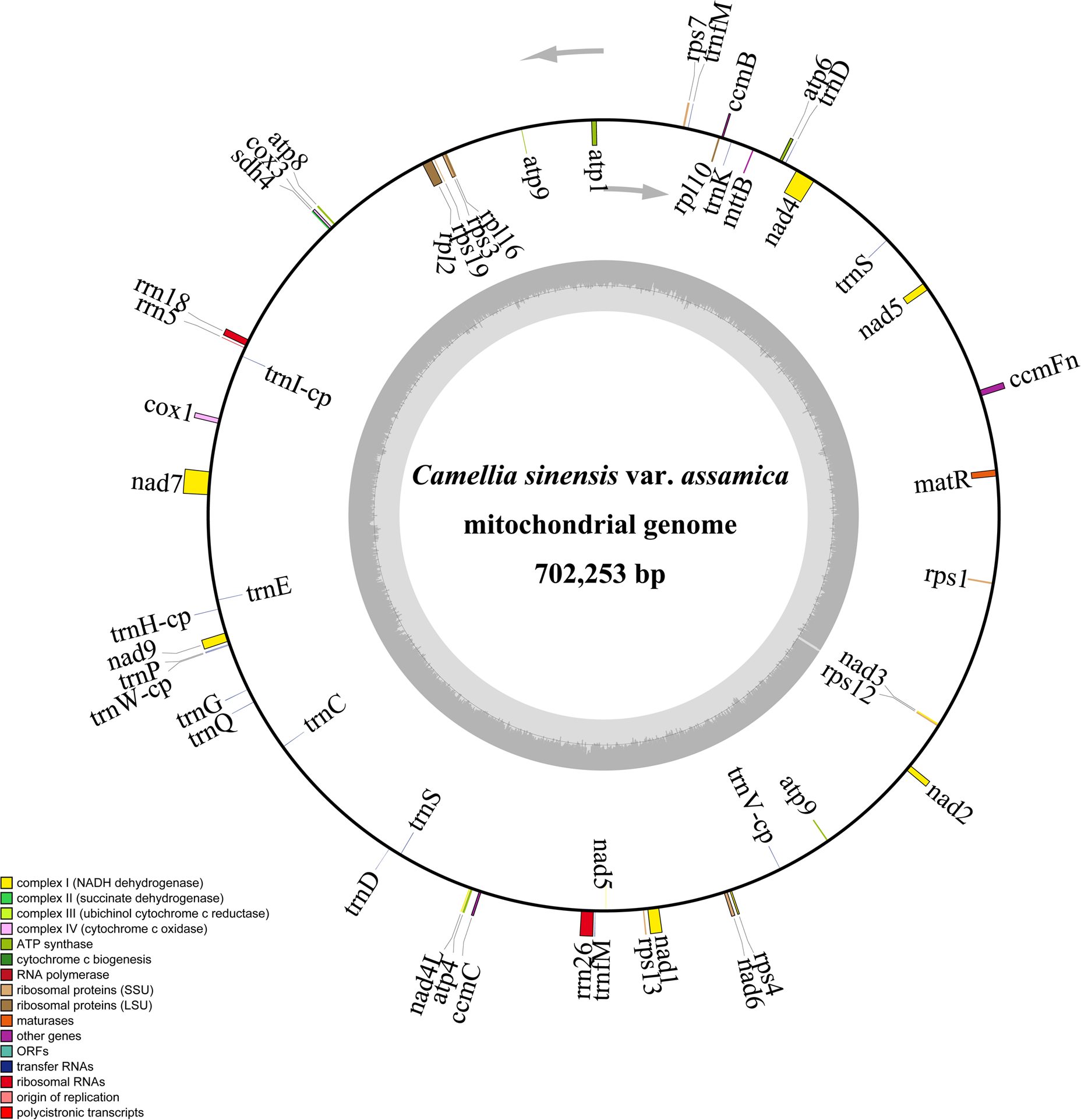
Circular map of scaffold 1 in the *C. sinensis* var. *assamica* cv. *Yunkang 10* mitochondrial genome. Gene map showing 54 annotated genes with different functional groups that are color-coded on outer circle as transcribed clock-wise (outside) and transcribed counter clock-wise (inside). The inner circle indicates the GC content as dark grey plot.

**Figure 4.**
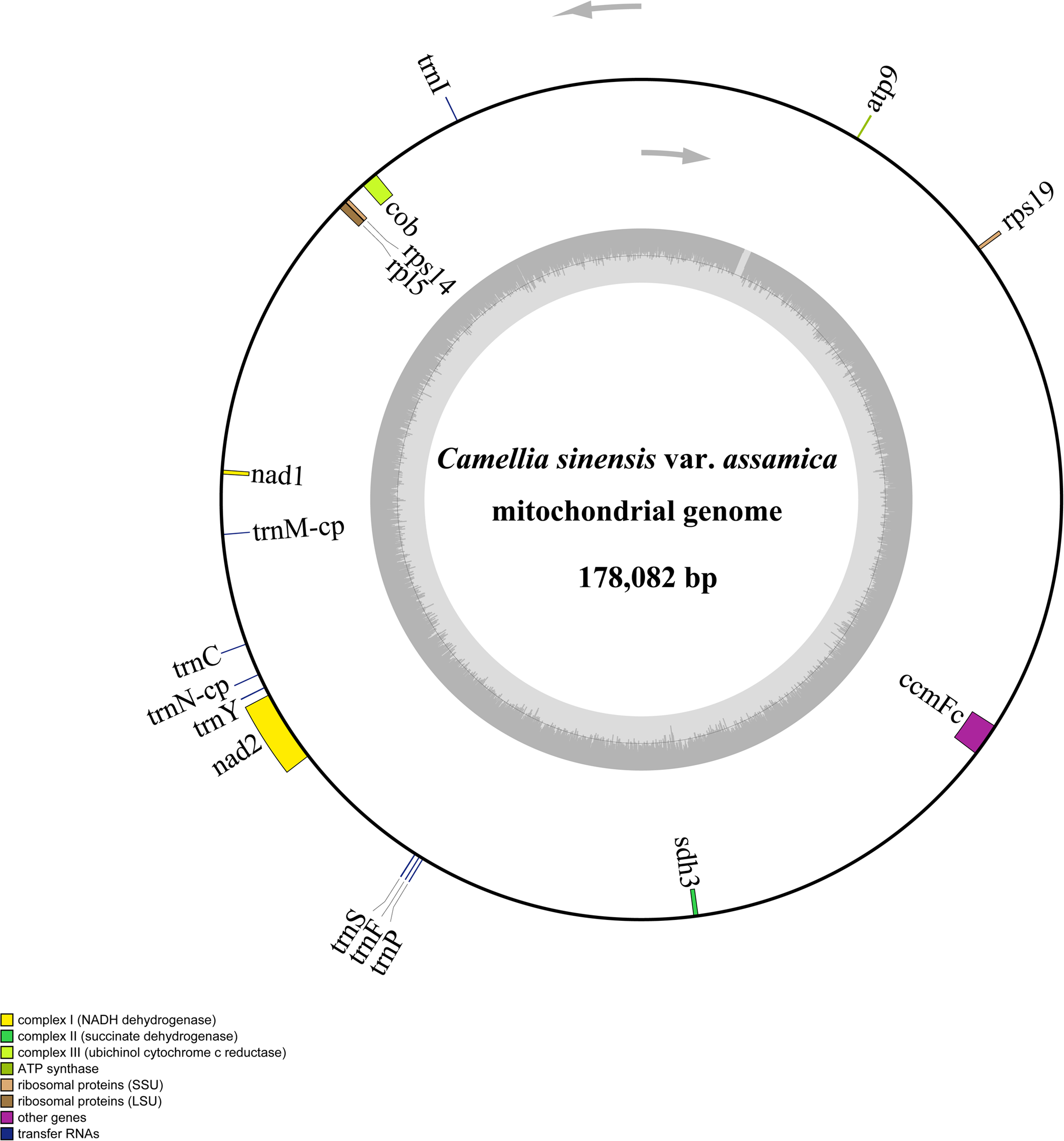
Circular map of scaffold 2 in the *C. sinensis* var. *assamica* cv. *Yunkang 10* mitochondrial genome. Gene map showing 17 annotated genes with different functional groups that are color-coded on outer circle as transcribed clock-wise (outside) and transcribed counter clock-wise (inside). The inner circle indicates the GC content as dark grey plot.

**Figure 5.**
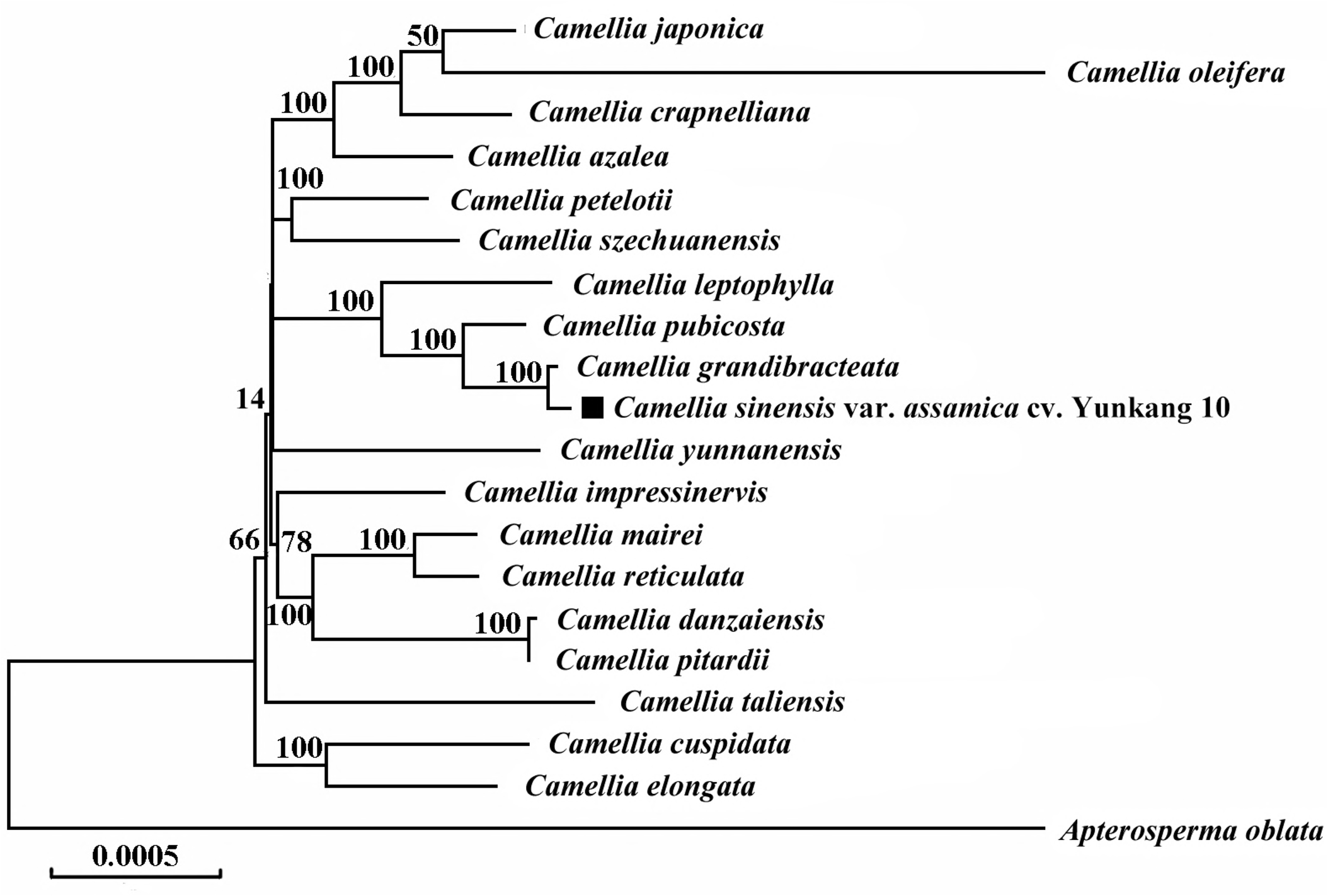
Phylogenetic relationships of 20 complete chloroplast genomes. Maximum likelihood phylogenetic tree of *C. sinensis* var. *assamica* cv. *Yunkang 10* with 18 species in the genus *Camellia* based on complete chloroplast genome sequences. The chloroplast sequence of *Apterosperma oblata* was set as outgroup. The position of *C. sinensis* var. *assamica* cv. *Yunkang 10* is shown in bold and bootstrap values are shown for each node.

**Figure 6.**
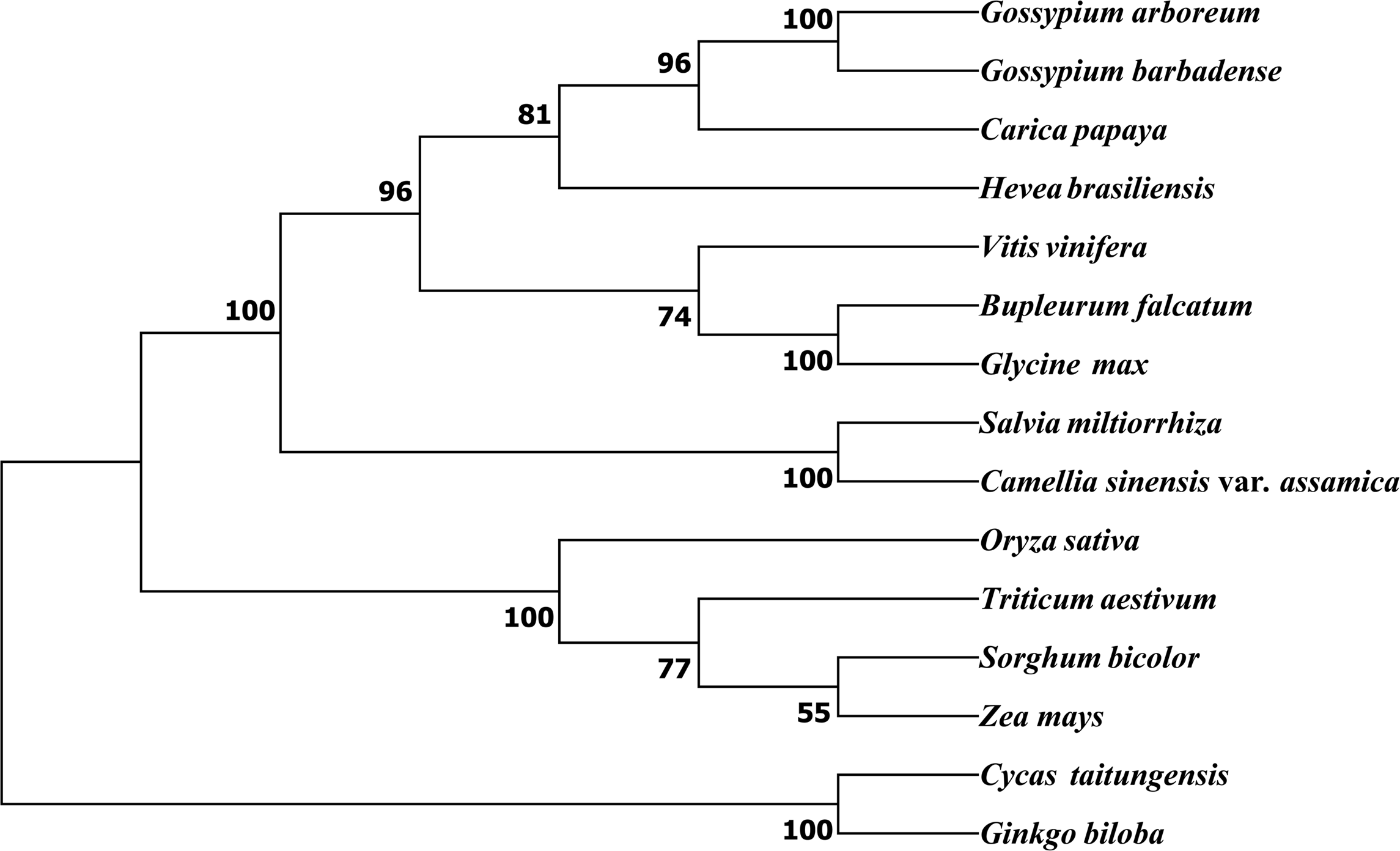
Phylogeny inferred from 13 genes common in the 15 plant mitochondrial genomes. Neighbor-joining tree of *C. sinensis* var. *assamica* cv. *Yunkang 10* with other 14 species based on 13 conserved protein-coding gene sequences with bootstrap support values on each node. The mt sequence of *Cycas taitungensis* and *Ginkgo biloba* were set as outgroup.

**Table 8.** Predicted RNA-editing sites in Scaffold 2 of the *C. sinensis* var. *assamica* mt genome*.

| No. | Gene | Nucleotide<br>Position | AA<br>Pos | Effect | Score |
| --- | --- | --- | --- | --- | --- |
| 1 | *rps19* | 116 | 39 | TCG (S) => TTG (L) | 1 |
| 2 | *rps19* | 163 | 55 | CCT (P) => TTT (F) | 1 |
| 3 | *rps19* | 164 | 55 | CCT (P) => TTT (F) | 1 |
| 4 | *atp9* | 53 | 18 | TCA (S) => TTA (L) | 1 |
| 5 | *atp9* | 83 | 28 | TCA (S) => TTA (L) | 1 |
| 6 | *cob* | 118 | 40 | CCG (P) => TCG (S) | 0.92 |
| 7 | *cob* | 178 | 60 | CAC (H) => TAC (Y) | 1 |
| 8 | *cob* | 286 | 96 | CTC (L) => TTC (F) | 1 |
| 9 | *cob* | 298 | 100 | CAC (H) => TAC (Y) | 1 |
| 10 | *cob* | 325 | 109 | CAT (H) => TAT (Y) | 1 |
| 11 | *cob* | 358 | 120 | CGG (R) => TGG (W) | 1 |
| 12 | *cob* | 419 | 140 | CCA (P) => CTA (L) | 1 |
| 13 | *cob* | 568 | 190 | CAT (H) => TAT (Y) | 0.92 |
| 14 | *cob* | 680 | 227 | TCT (S) => TTT (F) | 1 |
| 15 | *cob* | 808 | 270 | CCC (P) => TCC (S) | 1 |
| 16 | *cob* | 853 | 285 | CAT (H) => TAT (Y) | 1 |
| 17 | *cob* | 908 | 303 | CCA (P) => CTA (L) | 1 |
| 18 | *cob* | 914 | 305 | TCT (S) => TTT (F) | 1 |
| 19 | *cob* | 982 | 328 | CAC (H) => TAC (Y) | 0.85 |
| 20 | *cob* | 1015 | 339 | CGC (R) => TGC (C) | 1 |
| 21 | *cob* | 1084 | 362 | CCT (P) => TCT (S) | 1 |
| 22 | *cob* | 1124 | 375 | CCG (P) => CTG (L) | 1 |
| 23 | *rps14* | 47 | 16 | GCG (A) => GTG (V) | 0.6 |
| 24 | *rps14* | 271 | 91 | CCT (P) => TCT (S) | 0.6 |
| 25 | *rpl5* | 35 | 12 | TCA (S) => TTA (L) | 0.78 |
| 26 | *rpl5* | 47 | 16 | CCG (P) => CTG (L) | 1 |
| 27 | *rpl5* | 59 | 20 | CCG (P) => CTG (L) | 0.89 |
| 28 | *rpl5* | 64 | 22 | CAC (H) => TAC (Y) | 1 |
| 29 | *rpl5* | 92 | 31 | TCG (S) => TTG (L) | 1 |
| 30 | *rpl5* | 172 | 58 | CGC (R) => TGC (C) | 0.89 |
| 31 | *rpl5* | 518 | 173 | CCA (P) => CTA (L) | 0.89 |
| 32 | *rpl5* | 521 | 174 | CCG (P) => CTG (L) | 1 |
| 33 | *nad2* | 110 | 37 | TCT (S) => TTT (F) | 1 |
| 34 | *nad2* | 125 | 42 | TCC (S) => TTC (F) | 1 |
| 35 | *nad2* | 272 | 91 | TCT (S) => TTT (F) | 0.67 |
| 36 | *nad2* | 284 | 95 | TCA (S) => TTA (L) | 1 |
| 37 | *nad2* | 293 | 98 | TCT (S) => TTT (F) | 1 |
| 38 | *nad2* | 412 | 138 | CAT (H) => TAT (Y) | 1 |
| 39 | *nad2* | 442 | 148 | CGT (R) => TGT (C) | 0.78 |
| 40 | *nad2* | 446 | 149 | ACT (T) => ATT (I) | 1 |
| 41 | *nad2* | 512 | 171 | TCA (S) => TTA (L) | 0.78 |
| 42 | *nad2* | 542 | 181 | TCA (S) => TTA (L) | 1 |
| 43 | *nad2* | 611 | 204 | TCG (S) => TTG (L) | 1 |
| 44 | *nad2* | 731 | 244 | CCA (P) => CTA (L) | 0.67 |
| 45 | *nad2* | 760 | 254 | CGT (R) => TGT (C) | 1 |
| 46 | *nad2* | 932 | 311 | TCA (S) => TTA (L) | 0.67 |
| 47 | *nad2* | 941 | 314 | CCA (P) => CTA (L) | 1 |
| 48 | *nad2* | 989 | 330 | TCA (S) => TTA (L) | 1 |
| 49 | *sdh3* | 67 | 23 | CCA (P) => TCA (S) | 1 |
| 50 | *sdh3* | 74 | 25 | TCC (S) => TTC (F) | 1 |
| 51 | *ccmFc* | 38 | 13 | TCC (S) => TTC (F) | 0.83 |
| 52 | *ccmFc* | 50 | 17 | CCT (P) => CTT (L) | 1 |
| 53 | *ccmFc* | 52 | 18 | CGT (R) => TGT (C) | 1 |
| 54 | *ccmFc* | 103 | 35 | CCC (P) => TCC (S) | 1 |
| 55 | *ccmFc* | 119 | 40 | TCT (S) => TTT (F) | 1 |
| 56 | *ccmFc* | 122 | 41 | TCC (S) => TTC (F) | 1 |
| 57 | *ccmFc* | 146 | 49 | CCT (P) => CTT (L) | 1 |
| 58 | *ccmFc* | 151 | 51 | CCT (P) => TCT (S) | 0.83 |
| 59 | *ccmFc* | 155 | 52 | TCA (S) => TTA (L) | 1 |
| 60 | *ccmFc* | 160 | 54 | CCT (P) => TCT (S) | 0.67 |
| 61 | *ccmFc* | 203 | 68 | ACG (T) => ATG (M) | 1 |
| 62 | *ccmFc* | 305 | 102 | TCA (S) => TTA (L) | 0.83 |
| 63 | *ccmFc* | 391 | 131 | CGT (R) => TGT (C) | 1 |
| 64 | *ccmFc* | 406 | 136 | CGT (R) => TGT (C) | 0.83 |
| 65 | *ccmFc* | 620 | 207 | GCG (A) => GTG (V) | 1 |
| 66 | *ccmFc* | 704 | 235 | GCT (A) => GTT (V) | 0.83 |
| 67 | *ccmFc* | 1100 | 367 | CCA (P) => CTA (L) | 1 |
| 68 | *ccmFc* | 1121 | 374 | TCG (S) => TTG (L) | 1 |
| 69 | *ccmFc* | 1276 | 426 | CGA (R) => TGA (X) | 1 |

\*The cutoff score (C-value) was set to 0.6.

## Supplementary Legends

**Supplementary Table 1.** List of the 15 plant mitochondrial genome sequences used for phylogenetic analysis.

| **Organism Name** | **GenBank Accession Number** |
| --- | --- |
| *Gossypium arboreum* | NC_035073.1/KR736342.1 |
| *Gossypium barbadense* | NC_028254.1/KP898249.1 |
| *Carica papaya* | NC_012116.1/EU431224.1 |
| *Hevea brasiliensis* | AP014526.1 |
| *Vitis vinifera* | NC_012119.1/FM179380.1 |
| *Bupleurum falcatum* | NC_035962.1/KX887330.1 |
| *Glycine max* | NC_020455.1/JX463295.1 |
| *Salvia miltiorrhiza* | NC_023209.1/KF177345.1 |
| *Oryza sativa* | NC_011033.1/BA000029.3 |
| *Triticum aestivum* | NC_036024.1/EU534409.1 |
| *Sorghum bicolor* | NC_008360.1/DQ984518.1 |
| *Zea mays* | NC_007982.1/AY506529.1 |
| *Cycas taitungensis* | NC_010303.1/AP009381.1 |
| *Ginkgo biloba* | NC_027976.1/KM672373.1 |

